# Bet hedging buffers budding yeast against environmental instability

**DOI:** 10.1101/2020.04.08.032904

**Authors:** Laura E. Bagamery, Quincey A. Justman, Ethan C. Garner, Andrew W. Murray

## Abstract

To grow and divide, cells must extract resources from dynamic and unpredictable environments. Organisms thus possess redundant metabolic pathways for distinct contexts. In budding yeast, ATP can be produced from carbon by mechanisms that prioritize either speed (fermentation) or yield (respiration). We find that in the absence of predictive cues, cells vary in their intrinsic ability to switch metabolic strategies from fermentation to respiration. We observe subpopulations of yeast cells which either rapidly adapt or enter a shock state characterized by deformation of many cellular structures, including mitochondria. This capacity to adapt is a bimodal and heritable state. We demonstrate that metabolic preparedness confers a fitness advantage during an environmental shift but is costly in a constant, high-glucose environment, and we observe natural variation in the frequency of prepared cells across wild yeast strains. These experiments suggest that bet-hedging has evolved in budding yeast.

## INTRODUCTION

Metabolic flexibility--the ability to perform functionally interchangeable anabolic and catabolic activities through distinct pathways--allows cells to modulate their metabolic behavior in response to external nutrient supplies, which often vary widely in quantity and quality. Budding yeast can use multiple metabolic strategies to derive energy from carbon: in the presence of high concentrations of their preferred carbon source, glucose, yeast rely on glycolysis and fermentation for the generation of ATP and biomass; cells perform respiration, a process with a higher ATP yield per unit sugar, when glucose is scarce or when they depend on a carbon source that can only be metabolized through oxidation (Johnston and Carlson, 1992; Zaman et al., 2008).

This metabolic flexibility is constrained by the dynamics of the shift between fermentative and respiratory states. The budding yeast *Saccharomyces cerevisiae* prefers to ferment, a bias that is enforced by catabolite repression, a complex and overlapping set of regulatory networks that repress the expression of gene products involved in respiration even in the presence of oxygen and carbon sources that can be oxidized (Kayikci and Nielsen, 2015). The slow reversal of catabolite repression can delay cell proliferation. In a batch culture of yeast grown in typical synthetic medium, the initial reliance on heavy fermentation and the later respiration of the residual ethanol result in two separate exponential phases of growth with a few hours of apparently stalled growth between the two, known as the diauxic lag (Figure 1A).

**Figure 1.**
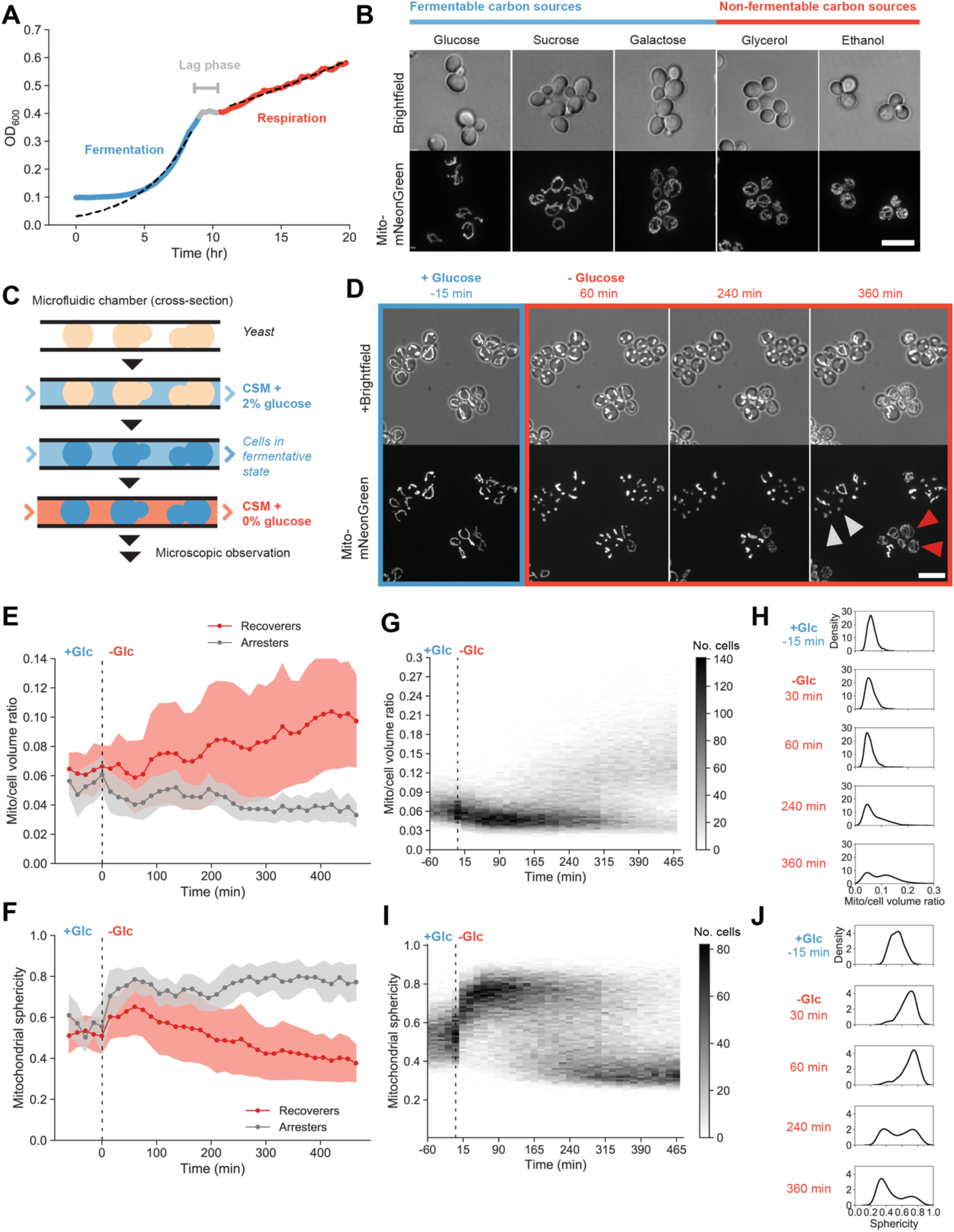
Heterogeneity in mitochondrial size scaling and structural integrity during fermentative-to-respiratory transitions. (A) Typical yeast growth dynamics, expressed as the optical density of a batch yeast culture (yLB126) measured on a plate reader with continuous shaking. Dotted lines depict exponential fits to the indicated blue and red regions. (B) Mitochondrial networks visualized by matrix-targeted mNeonGreen fluorescent protein (mito-mNeonGreen) (yLB126) in exponentially-growing yeast cells in synthetic media with indicated carbon sources. Scale bar, 10 μm. (C) Experimental setup. Cells were immobilized in microfluidic plates and supplied with synthetic media containing 2% glucose by continuous perfusion, promoting glucose repression and fermentative growth. Cells were observed by microscopy during this pregrowth period and following abrupt replacement of the source media with synthetic media containing 0% glucose. (D) Representative images of mito-mNeonGreen (yLB126) in high glucose and followed by abrupt switch to glucose-free media. Labeled cells indicate examples of recoverer cells, which adapt and perform mitochondrial biogenesis (red arrows), and arrester cells, with extended mitochondrial collapse (gray arrows). Scale bar, 10 μm. (E) Distributions of the ratio of mitochondrial to total cell volume for phenotypic classes above for all cells in the field of view in *(D)*. Filled bands indicate one standard deviation from the mean. These sample trajectories are representative of 13 independent experiments as displayed in *(G)* and *(H)*. (F) Distributions of mitochondrial sphericity index (ratio of surface area of hypothetical sphere with the same volume as a mitochondrion to its actual surface area) for phenotypic classes in *(D)*, in a given sample field of view. Filled bands indicate one standard deviation. (G) and (H) Time-resolved heat maps of mitochondrial/cell volume ratio *(G)* and mitochondrial sphericity *(H)* in N = 1,329 cells, collected across 13 independent experiments, before and during acute glucose starvation beginning at 0 min. Intensity reflects absolute number of cells within the binning area. (I) and (J) Histograms of mitochondrial/cell volume ratio *(I)* and sphericity *(J)* at the indicated time points from *(G)* and *(H)* displaying bimodality in mitochondrial morphology following glucose withdrawal. See also Figure S1.

To understand the nature of the lag phase and metabolic transitions more generally, we examined mitochondria as markers of metabolic state in individual cells undergoing nutrient shifts. In yeast, mitochondria exist as a network of branched tubules along the cell cortex. Mitochondrial network size, structure, and ultrastructure are flexible and responsive to environmental conditions: cells in the presence of high glucose possess minimal mitochondrial networks that can increase in both volume and the number of branching points when forced to utilize a nonfermentable carbon source such as glycerol (Egner et al., 2002; Hoffmann and Avers, 1973; Stevens, 1977; Yotsuyanagi, 1962).

We examined the dynamics of fermentative-to-respiratory shifts in single cells by using microfluidics to rapidly switch between different carbon sources. We measured mitochondrial network size and morphology as markers of cellular reorganization following abrupt glucose deprivation. We observed heterogeneity across isogenic populations: mitochondrial networks lose structural integrity and collapse into spherical globules in most cells, but one population recovers a tubular morphology and resumes proliferation while the other remains in a state of arrest. This difference was accompanied by larger-scale organellular perturbation and was heritable. We identified a set of mutations that display homogeneous recovery upon acute glucose starvation, all of which directly affect glucose sensing, signaling, or utilization. These strains possess larger and more branched mitochondrial networks and partly depend on respiration even when grown on glucose. Growth on non-fermentable carbon sources selects for cells capable of recovery, but in the presence of glucose, cells rapidly switch into the arrester state while the reverse transition occurs more slowly, such that arrester cells predominate after prolonged growth on glucose. Analyzing the growth of single cells reveals that capacity for recovery imposes a short-term fitness cost. This environmental growth-rate tradeoff also occurs in natural yeast strains, and their different degrees of metabolic flexibility suggest that the extent of preparation against nutrient instability is under evolutionary selection.

## RESULTS

### Bimodality in mitochondrial structural integrity and organelle size scaling during sudden glucose deprivation

To assess whether mitochondrial network morphology reports on a yeast cell’s metabolic state, we examined mitochondria in cells growing in a variety of carbon sources. To visualize mitochondria, we fused a mitochondrial targeting sequence to the fluorescent protein mNeonGreen (mito-mNeonGreen), which we expressed constitutively. The gene product is transported to the mitochondrial matrix and the targeting sequence cleaved off, labeling the mitochondrial matrix with free fluorescent protein.

As described previously (Egner et al., 2002; Stevens, 1977; Yotsuyanagi, 1962), microscopic images reveal divergent mitochondrial network architectures as a function of nutrient availability, with minimal mitochondrial content in the presence of high levels of fermentable sugars (glucose, sucrose, galactose) but with denser, more extensive, mitochondrial networks in cells growing under respiratory conditions (glycerol, ethanol) (Figure 1B). We performed three-dimensional segmentation of mitochondrial networks and calculated the proportion of the total cell volume occupied by mitochondrial content. We found that distinct carbon sources produced mitochondrial volume ratios spanning a nearly two-fold range, from 0.06 under high glucose conditions to 0.11 in cells grown on ethanol (Figure S1A). To quantify the relationship between network size and metabolic output, we measured oxygen consumption rates as a proxy for respiratory activity. Cells with larger mitochondrial networks depleted oxygen from the medium faster, indicating that increased respiratory activity correlates with increased mitochondrial content (Figure S1B).

We then followed mitochondria during metabolic transitions by fluorescent microscopy. We used microfluidics to rapidly switch the media perfusing immobilized cells (Figure 1C). After an initial flow of synthetic media containing high glucose to promote fermentation, cells were shifted into glucose-free medium to prevent fermentation but allow the respiration of amino acids in the medium.

We observed a heterogeneous response to sudden glucose deprivation (Figure 1D and Movies S1-S2). All cells arrested their growth at the moment of the loss of glucose from the medium. In most cells, mitochondrial networks lost their tubular structure and condensed into spherical globules. In a subset of these cells, this collapse in mitochondrial structural integrity suddenly and rapidly reversed itself at varying time intervals and was followed by an increase in mitochondrial network volume and the eventual resumption of growth. The rest remained arrested indefinitely, though such cells remained physiologically and genetically viable: they recovered their mitochondrial structure and resumed growth and division when glucose was returned to the environment (Movie S2). We observed a plateau at roughly 6 hr post-starvation after which cells ceased to transition from the arrested to the recovering state, and these states’ relative proportions remained stable for at least another 4 hr (Figure S1C).

We quantified the dynamics of these morphological and volumetric changes in the mitochondrial network. Figure 1E displays mitochondrial trajectories for the cells shown in Figure 1D, classified as recovering or arresting based on their possession of tubular or spherical mitochondria, respectively, at 6 hr post-starvation. We measured the total mitochondrial volume within these cells, expressed as a proportion of total cell volume, and observed an approximate doubling of mitochondrial network size in cells whose mitochondria were tubular.

To assess the tubularity of the network, we measured the sphericity index, a dimensionless metric for divergence from the proportions of a sphere (see Methods). We observed increases in mitochondrial sphericity beginning at the time glucose was removed and peaking approximately 1 hr into starvation (Figure 1F). In some cells, this value remained constant over time. In others, sphericity abruptly decreased to a new steady-state lower than the initial pre-starvation value, reflective of both the reestablishment of mitochondrial tubularity and mitochondrial biogenesis, as a larger mitochondrial volume is arranged into longer rodlike structures with more branching points within the same, limited cell volume. This feature of mitochondrial sphericity allows us to distinguish larger-scale losses of structural integrity, seen as dramatic increases in the sphericity index, from remodeling activity associated with the establishment of new mitochondrial network sizes, detected in subtler decreases from the baseline.

We present the heterogeneity in starvation responses on a population scale in Figure 1G. We observe initial mitochondrial-to-total-cell volume ratios near 0.06, with an asynchronous rise to post-starvation to ratios centered about 0.13 as well as a subpopulation that does not increase its mitochondrial volume fraction (Figure 1G, with selected histograms in Figure 1H). Similarly, mitochondrial sphericity indices, initially concentrated between 0.4 and 0.6, rise to 0.8, then decrease in a subset of cells to values centered about 0.35 (Figures 1I and 1J). Histograms plotted from individual time points demonstrate the bimodality within the distributions of both mitochondrial volume ratio and sphericity, with post-starvation cells concentrated about two distinct peaks reflecting the presence or absence of mitochondrial biogenesis and the retention or loss of structural integrity (Figures 1H and 1I).

### Bimodal effects on growth, pH homeostasis and other cellular structures during environmental instability

The aim of metabolic adaptation is to support growth and division cycles in the altered environment. To determine the relationship between mitochondrial structural properties and proliferative capacity, we determined the time of first visible cell volume increase (or bud emergence) for a set of 357 cells from the recoverer subpopulation and measured their mitochondrial networks at this transition point. Relative to the predominantly collapsed mitochondrial state observed at 1 hr post-starvation and the bimodal distribution of collapsed and recovered cells at 6 hr starvation, the distribution of cells just beginning regrowth is a unimodal distribution of low sphericity, matching one of the two subpopulations within the larger data set (Figure 2A). Likewise, relative to the consistently low mitochondrial volume ratio in cells cultivated in the presence of high glucose and the mixture of low and high ratios present 6 hr post-starvation, the mitochondrial volume ratios of growing cells in glucose-deprived conditions are uniformly high and overlap with the upper peak of the bimodal distribution (Figure 2B). We observed no growth in cells whose mitochondria did not recover from their initial collapse. Cells that did recover had restored mitochondrial tubularity and increased mitochondrial volume more than 90 min prior to resuming growth (Figures S2A and S2B). We conclude that cells in a state of mitochondrial collapse are nongrowing cells which have not begun preparatory functions for respiration, including mitochondrial biogenesis.

**Figure 2.**
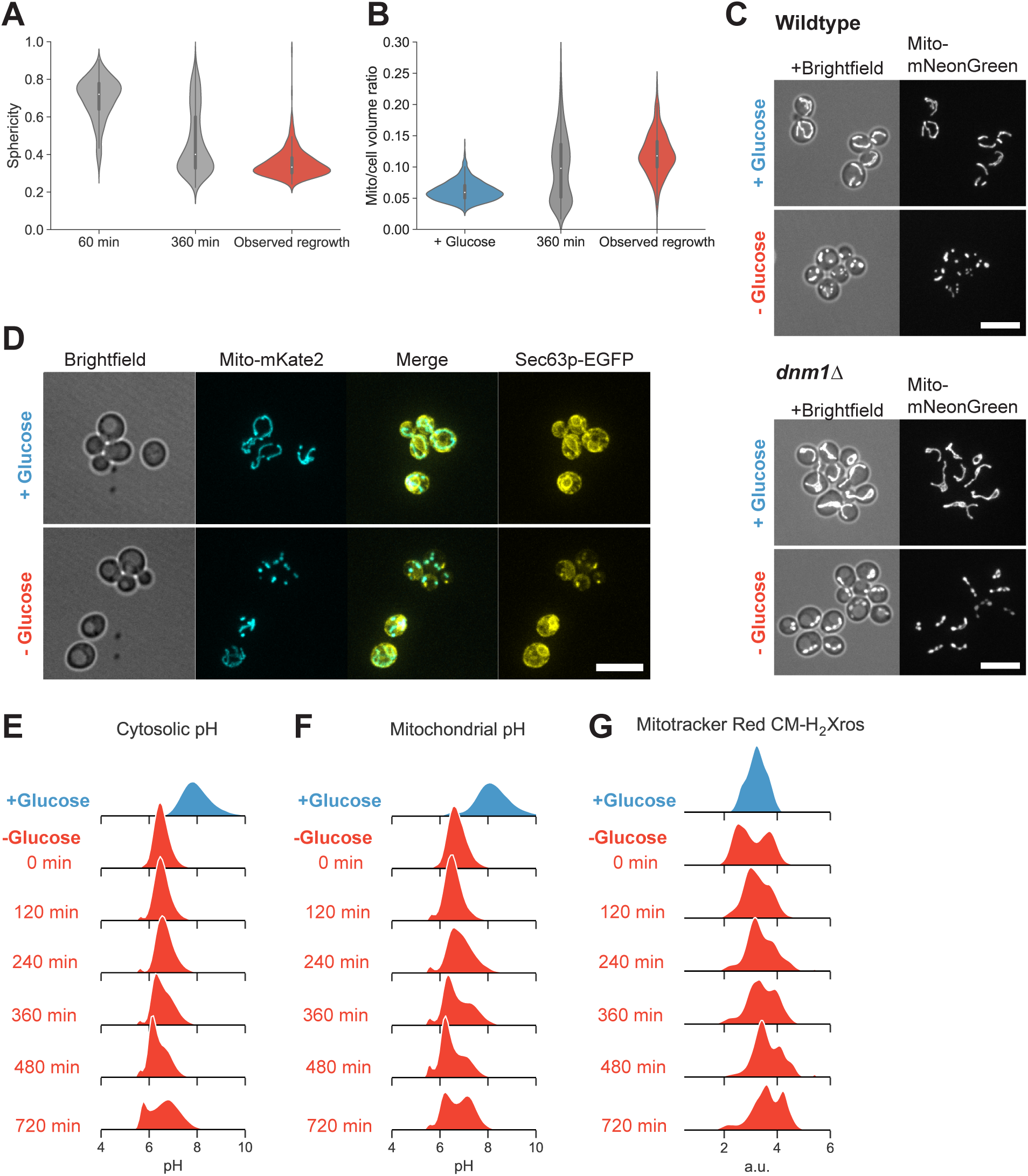
Mitochondrial structural collapse is accompanied by internal pH drop and other intracellular aggregation. (A) Distributions of mitochondrial sphericity index following 1 and 6 hr acute glucose starvation (N ≥ 357) and in a subpopulation of recovering cells at the time they resumed growth as determined by observable cell volume increase and/or budding. (B) Distributions of mitochondrial to total cell volume ratio in cells growing in synthetic media containing high glucose, cells starved of glucose for 6 hr, and starving cells at the first detectable resumption of growth. (C) Mitochondrial matrix marker in wild-type (yLB126) and mitochondrial fission-defective *dnm1Δ* mutants (yLB134), in synthetic media containing high glucose and following abrupt glucose withdrawal. Scale bars, 10 μm. (D) Co-expression of mito-mNeptune and the endoplasmic reticulum marker Sec63p-mNeonGreen (yLB41) during exponential growth in synthetic media containing high glucose and following glucose washout. Scale bar, 10 μm. (E)-(F) Dynamics of intracellular pH *(E)* and mitochondrial pH *(F)* in cells growing in high glucose and post-starvation, detected by constitutively-expressed pHluorin2 localized to the cytosol or mitochondrial matrix, respectively (yLB397 and yLB219). (G) Relative potential across the inner mitochondrial membrane before and following glucose starvation, measured in cells (yLB1) stained with potential-sensitive MitoTracker Red CM-H2-Xros. All distributions consist of three biological replicates of n = 40,000 cells. See also Figure S2.

The disappearance of tubular mitochondria could stem either from unbalanced mitochondrial fission, producing unusually short tubules that could aggregate into larger spheres, or by inhibition of the (largely uncharacterized) mechanisms that maintain mitochondrial rodlike structure, thereby allowing individual mitochondrial to balloon from tubules into globules. We examined the role of mitochondrial fission in these morphological changes by examining starvation dynamics in a fission-deficient *dnm1*Δ mutant. When growing in the presence of high glucose, *dnm1*Δ cells possess a single mitochondrial network with extensive branching; once glucose is removed from the media, the continuity of the network is maintained, but multiple structural collapses occur, resulting in a series of globules connected by small bridges (Figure 2C). From this, we conclude that mitochondrial fission is not the mechanism of collapse.

To determine the extent to which loss of mitochondrial structural integrity may instead reflect higher-order stress, we examined other cellular structures. Given the dependence of mitochondrial tubularity on mitochondrial contact with endoplasmic reticulum (Burgess et al., 1994; Sogo and Yaffe, 1994), we examined mitochondria in tandem with endoplasmic reticulum structure via a fusion of the fluorescent protein EGFP to the endoplasmic reticulum membrane protein Sec63p (Deshaies et al., 1991; Toyn et al., 1988). We found that arrested cells experiencing mitochondrial collapse during acute glucose starvation also contained aggregates of Sec63p which were absent in recoverer cells with tubular mitochondria (Figure 2D). We also observed aberrant localization patterns of the actin filament-binding protein Abp140p (Figure S2D) and the cortical actin patch protein Abp1p (Figure S2E), again only in arrester cells with mitochondrial collapse.

Previous studies have observed large protein assemblies and aggregation associated with nutrient starvation in budding yeast (Narayanaswamy et al., 2009; Petrovska et al., 2014; Suresh et al., 2015; Zacharogianni et al., 2014), postulated to be a consequence of cytosolic acidification resulting from the absence of glucose metabolism and a concurrent inability to maintain pH homeostasis (Dechant et al., 2010; Munder et al., 2016; Orij et al., 2009). We thus measured both the cytosolic and mitochondrial pH of cells induced to switch metabolic states by acute glucose starvation.

We measured pH using the pHluorin2, a fluorescent protein with a ratiometric signal that is pH-sensitive (Mahon, 2011). We opted to measure pH dynamics by flow cytometry, in samples arrested in bulk culture by centrifugation and resuspension in media lacking glucose (see Methods). Measurements of pHluorin2 are sensitive to noise from two intensity measurements, and one excitation wavelength, 395 nm, is particularly phototoxic to yeast, disfavoring longer-term live imaging.

Consistent with previous literature, we found that cytosolic pH rapidly dropped upon glucose depletion, from a mean of nearly pH 8 to 6.5 (Figure 2E). Upon prolonged starvation, however, we observed a bimodal pH distribution centered about pH 6 and 6.7, with one subpopulation recovering slightly from the initial acidification event, reminiscent of the heterogeneity observed in mitochondrial collapse and partial recovery. We observed a subtle but consistent cytosolic pH decrease in cells containing collapsed mitochondria relative to cells retaining tubular structure when we examined starving populations by microscopy, confirming that the subset of cells with disrupted pH homeostasis are those with similarly perturbed mitochondrial morphology (Figure S2E). We also observed a dramatic pH drop inside the mitochondrial matrix followed by eventual development of a bimodal distribution as only a portion of cells manage to increase matrix pH thereafter (Figure 2F).

We applied a potential-sensitive fluorescent mitochondrial dye, MitoTracker Red CM-H_2_Xros, to cells before and following starvation to determine whether there is a corresponding disruption of the electrical potential across the inner mitochondrial membrane. This potential, in conjunction with the pH gradient created by the expulsion of protons from the matrix, provides the free energy gradient necessary to drive protons through the ATP synthase complex and generate ATP, the prime mitochondrial respiratory function (Nicholls, 2002, 2004). We observed bimodality in mitochondrial potential immediately post-starvation, with a subpopulation of cells experiencing a steep loss of potential, followed by a continued two-peaked distribution in which the worst-affected cells partially increased their membrane potential but remained consistently distinguishable from the second subpopulation (Figure 2G).

A sudden starvation event thus produces a population that is heterogeneous not just in mitochondrial structure but in the integrity of a variety of cellular structures and processes, such that some cells successfully adapt and resume growth while others experience large-scale disruption of mitochondrial tubularity, actin networks, endoplasmic reticulum morphology, cytosolic and mitochondrial pH, and mitochondrial membrane potential.

### Heterogeneity during nutrient transitions is a heritable trait specific to carbon sugar metabolism

To determine whether this unusual mitochondrial stress response and the variable capacity for recovery are features of a general starvation response or unique to carbon source starvation, we examined mitochondrial dynamics during the abrupt disappearance of another nutrient, nitrogen, from the extracellular environment. We cultured prototrophic cells, capable of growing without external amino acid supplements, in minimal synthetic medium containing ammonium sulfate as the sole nitrogen source and glucose as the sole carbon source and performed media switching to deprive cells of either extracellular nitrogen or carbon.

As in previous experiments, removing glucose led to the collapse of mitochondrial network structure and recovery--or retention of tubularity--followed by network size increase in only a subpopulation of cells (Figure 3A, left); unlike their behavior in complete media, recoverer cells with tubular mitochondria never resumed the cell cycle or produced new buds, indicating that cells adapting to the absence of carbon utilize amino acids as an alternative carbon source (Figure S3A). However, cells deprived of nitrogen displayed neither the mitochondrial collapse nor the subsequent mitochondrial biogenesis that occurs in recovering cells after carbon starvation (Figures 3A-B and S3B).

**Figure 3.**
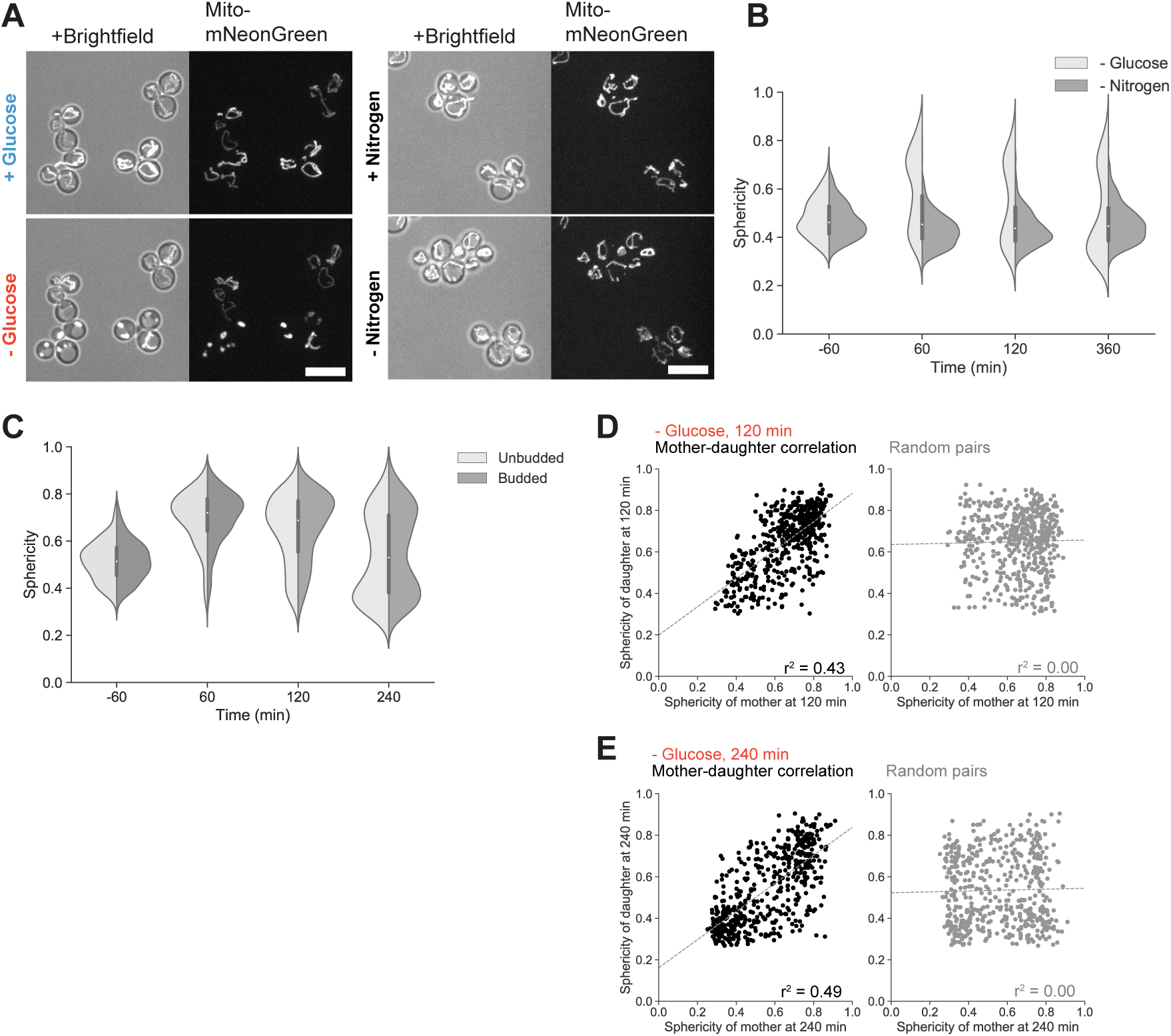
Glucose specificity and heritability of mitochondrial structural collapse. (A) Mitochondrial matrix fluorescent marker in prototrophic cells growing in synthetic media lacking amino acids, in the presence of high glucose and 1 hr following glucose withdrawal (left), or in the same synthetic media, before and 1 hr following nitrogen (ammonium sulfate) withdrawal (right panel). Scale bars, 10 μm. (B) Distribution of mitochondrial sphericity in cells that were acutely deprived of glucose or nitrogen at time 0 min, N ≥ 360 cells. (C) Mitochondrial sphericity in cells before and following glucose washout at time 0 min, partitioned by budded or unbudded state at the moment of glucose withdrawal. N ≥ 404 cells per type. (D)-(E). Correlation between mitochondrial sphericity in mother-daughter pairs following starvation for 120 min *(D)* and 240 min *(E)*, contrasted with correlation between mother and daughter cells when paired at random. Dotted lines depict regression fits calculated by least-squares method. N = 617 mother-daughter cell pairs. See also Figure S3.

We wondered whether this mitochondrial response stemmed from the loss of an easily metabolizable carbon source rather than another stress, such as sudden osmotic shock, that might accompany glucose disappearance. We performed a similar carbon source replacement in which glucose withdrawal was accompanied by its simultaneous replacement with galactose, a fermentable carbon source that nevertheless requires additional enzymes, normally repressed in the presence of glucose, for its conversion into glucose-6-phosphate (Timson, 2007). We again observed mitochondrial collapse and bimodality in mitochondrial structure, even in the presence of a fermentable carbon source (Figure S3C), suggesting that this effect is specific to the loss of glucose and not its indirect effects on extracellular conditions.

We wished to understand how a genetically identical population could display two such divergent phenotypes upon sudden glucose starvation. The most obvious source of non-genetic heterogeneity is cell cycle state, as budding and division cycles typically proceed asynchronously within a yeast population. We examined whether differential sensitivity to starvation and subsequent growth capacity could be explained by cell cycle heterogeneity by partitioning our data on the basis of each individual cell’s budded or unbudded status at the moment of disappearance of glucose from the media. We found that each population displayed a dramatic initial increase in mitochondrial sphericity followed by the development of a bimodal distribution of cells, eliminating cell cycle stage as a determinant of post-starvation behavior (Figures 3C and S3D).

We next examined whether the response to abrupt glucose deprivation was instead driven by lineage-dependent heterogeneity. We identified pairs of mother and daughter cells and compared their responses to glucose withdrawal. We found that cells and their direct progenitors were likely to similarly suffer from or avoid mitochondrial collapse, with specific sphericity indices modestly correlated (r^2^ = 0.43) relative to randomly paired mothers and daughters (r^2^ = 0.00) after 2 hr starvation (Figure 3D). This lineage correlation was maintained as the adapting subpopulation began to alter its sphericity distribution 4 hr post-starvation (mother-daughter r^2^ = 0.49; random pairing r^2^ = 0.00) (Figure 3E).

### Glucose sensing, signaling, and utilization pathways modulate heterogeneity in cell starvation fates

To elucidate the factors influencing cell fate during acute starvation, we searched for mutants with unimodal starvation responses, focusing on the loss of gene products involved in glycolysis and associated regulatory activity. We identified four sets of single and double deletions that permit cells to recover uniformly and rapidly during sudden starvation by variously disrupting external glucose sensing (*rgt2*Δ *snf3*Δ double mutant), glucose phosphorylation (*hxk2*Δ), glucose-dependent transcriptional repression (*mig1*Δ *mig2*Δ double mutant), or the ability to inhibit the Snf1p kinase complex, a crucial positive regulator of respiratory induction (*reg1*Δ) (Figure 4A). The opposite phenotype, uniform and indefinite loss of mitochondrial structural integrity, is associated with loss of Snf1p kinase itself (*snf1*Δ). Representative images of homogeneous mitochondrial collapse or recovery and adaptation are shown in Figure 4B and their sphericity dynamics quantified in Figure 4C (see also Movie S3). The mutants which prevent mitochondrial collapse do not display detectable increases in sphericity upon sudden starvation. In contrast, *snf1*Δ mutants undergo similar structural collapse as wild-type cells but never develop a subpopulation of recovering cells (see also Figures S4A and S4B).

**Figure 4.**
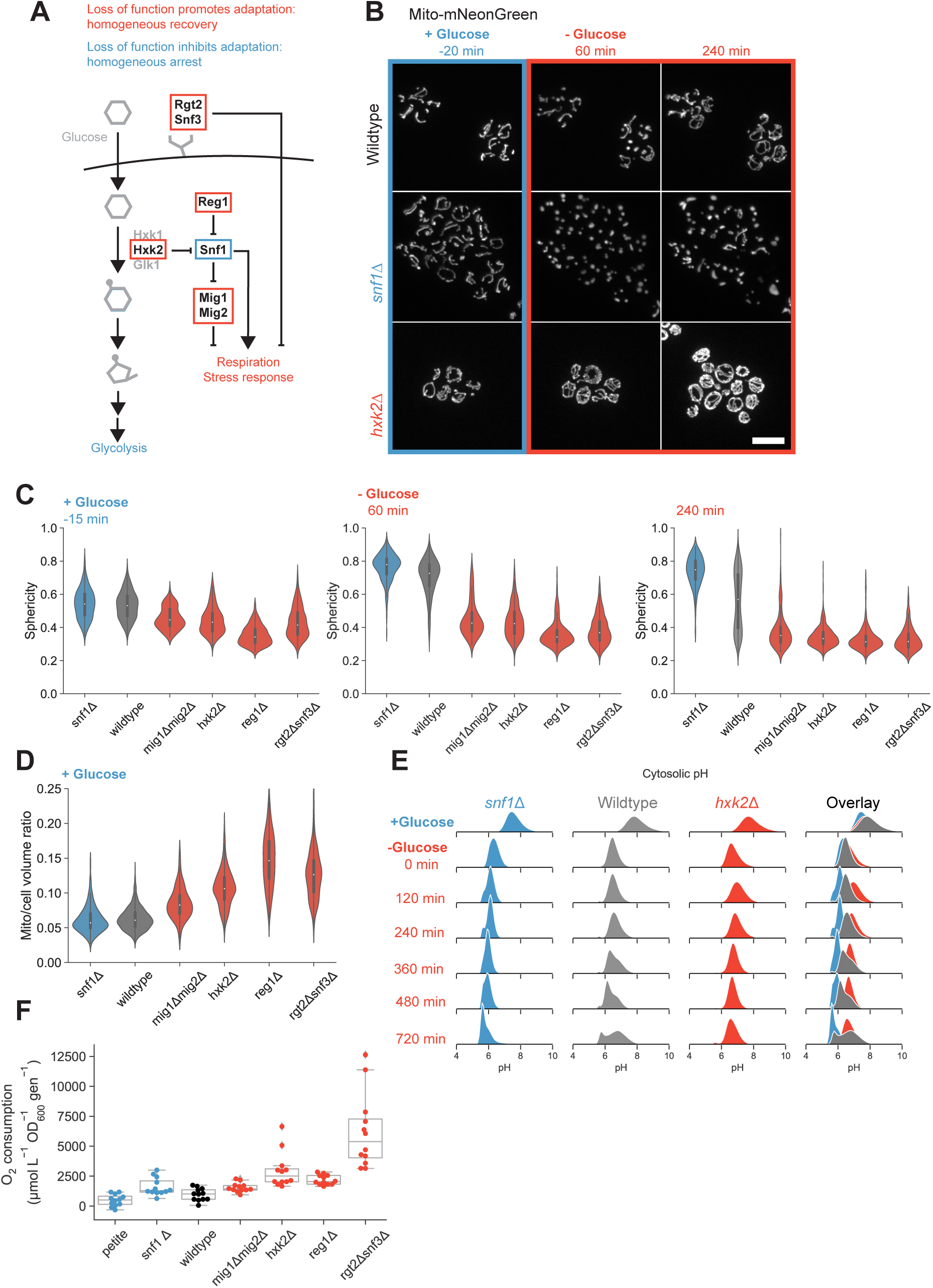
Glucose signaling and utilization pathways modulate post-starvation behavior. (A) Simplified depiction of selected glucose sensing and signaling pathways in budding yeast. Red boxes indicate gene products for which loss of function results in rapid, homogeneous adaptation during acute glucose starvation. The blue box identifies a gene product whose loss results in a homogeneous inability to adapt following glucose withdrawal. (B) Mitochondrial matrix marker in wild-type (yLB126), *hxk2*Δ (yLB146), and *snf1*Δ (yLB168) cells, growing in synthetic media containing high glucose and following abrupt removal of glucose by washout. Scale bar, 10 μm. (C) Distribution of mitochondrial sphericity in *snf1*Δ (yLB168), wild-type (yLB126), *mig1*Δ*mig2*Δ (yLB180), *hxk2*Δ (yLB146), *reg1*Δ (yLB196), and *rgt2*Δ*snf3*Δ (yLB233) strains growing in high glucose (left panel) and 60 min and 240 min post-glucose withdrawal. Mutants in red display relatively low sphericity and no increase post-starvation. The *snf1*Δ mutant in blue initially resembles wild-type but uniformly fails to adapt during starvation. N ≥ 142 cells per genotype. (D) Pre-starvation mitochondrial to cell volume ratios for the strains in *(C)*. N ≥ 142 cells per genotype. (E) Cytosolic pH in wild-type (yLB397), *hxk2*Δ (yLB416), and *snf1*Δ (yLB412) cells expressing ratiometric pHluorin2, prior to and following glucose starvation. All distributions consist of three biological replicates of N = 40,000 cells each. (F) Oxygen consumption rates, normalized by growth rate, for petite (yLB373), *snf1*Δ (yLB167), wild-type (yLB1), *mig1*Δ*mig2*Δ (yLB181), *hxk2*Δ (yLB146), *reg1*Δ (yLB194), and *rgt2*Δ*snf3*Δ (yLB232) strains growing in synthetic media containing high glucose. Data consist of three biological replicates, each comprised of four technical replicates. All Kolmogorov-Smirnov test p-values against wild-type < 0.05 save for petite mutant (p = 0.19). See also Figure S4.

The lower sphericity indices in our suppressor strains even in the presence of abundant glucose suggested that their mitochondrial networks may be larger prior to any nutrient shift, which we confirmed by calculating mitochondrial to total cell volume ratios ranging from 0.092 (*mig1*Δ*mig2*Δ) to 0.147 (*reg1*Δ), in contrast to ratios of 0.061 and 0.063 in wild-type and *snf1*Δ cells, respectively (Figure 4D).

We wondered whether the homogeneity in mitochondrial fates that we observed in association with these mutants extended to other intracellular manifestations of stress, particularly the extreme pH drop observed immediately following glucose washout. The mean cytosolic pH of wild-type, *hxk2*Δ, and *snf1*Δ cells expressing pHluorin2 is initially similar, at roughly pH 7.6, and all three strains experience cytosolic acidification upon loss of glucose (Figure 4E). However, as wild-type cells partially adapt to produce a two-peaked distribution, the pH of each mutant is unimodal and overlaps with one of the two wild-type subpopulations, *snf1*Δ cells retaining a low pH and *hxk2*Δ mutants reaching a new steady-state of slightly increased pH several hours in advance of the recovering wild-type cells. Each of these mutant strains thus appears to recapitulate one of the two wild-type subpopulations, albeit with faster recovery dynamics in the case of *hxk2*Δ.

Even when they are grown on high glucose, our collapse-suppressing, fast-adapting mutants are all closer in mitochondrial content to glucose-starved cells than they are to wild-type cells. We therefore hypothesized that the mechanism by which these mutations permit homogeneous recovery was that they were prepared for starvation: they had already prepared for or adopted a respiratory metabolic state before glucose disappeared from the media without warning. Prior work has demonstrated that a history of growth on a non-glucose carbon source can decrease lag time during subsequent shifts from glucose to other non-glucose sugars (Cerulus et al., 2018).

To test whether the collapse suppressors were respiring in glucose, we measured oxygen consumption rate as a reporter of respiratory activity in our wild-type cells, five mutant strains, and a respiratory-deficient petite strain growing in the presence of high glucose (Figure 4F). We found that wild-type cells displayed a rate of oxygen depletion and respiration that did not differ significantly from that of petites (p = 0.19). The consumption rates of all four suppressor/recoverer mutants were in turn significantly elevated above wild-type (p ≤ 0.02). Curiously, *snf1*Δ mutants, which are also respiratory-defective, consumed oxygen more rapidly than our wild-type strain (p = 0.02). This finding is in agreement with previous work which has detected increased oxygen consumption by *snf1*Δ mutants in media containing 2% glucose, postulated as an increase in oxidative phosphorylation using amino acids from the media to partially compensate for lack of Snf1p function while proving insufficient in scale for ATP generation or cell survival on purely respiratory carbon sources (Nicastro et al., 2015). With the exception of this anomalous oxygen consumption by respiratory-defective *snf1*∆ mutants, whose physiological purpose remains an open question in yeast metabolism, these data are consistent with the notion that prior respiratory activity is beneficial during sudden starvation events, and we observe no instances in which any of our suppressors adapt during acute starvation without some degree of constitutive respiration.

### Preparation for future starvation allows for faster adaptation at an immediate fitness cost

We wondered whether the relationship that we observed between efficient starvation recovery and initial mitochondrial network size could explain the variability in wild-type populations. We ranked our data set of 1,329 cells by their mitochondrial to total cell volume ratios at the time immediately before abrupt glucose deprivation and tracked the least- and most-mitochondrially endowed 10% of cells across the environmental transition. We observed a bias towards rapid recovery, often within one hour, in our high-mitochondrial subgroup relative to the larger population, as well as an enrichment in recoverers more generally (Figures 5A and 5C). In contrast, cells with the lowest mitochondrial volume fraction in the presence of high glucose were less likely to decrease in sphericity after the initial structural shock, and the small percentage of cells that did not observably increase in sphericity at all were absent from this subpopulation (Figures 5B and 5D). The proportion of the cell’s volume allotted to the mitochondrial network thus has predictive value for a cell’s future behavior.

**Figure 5.**
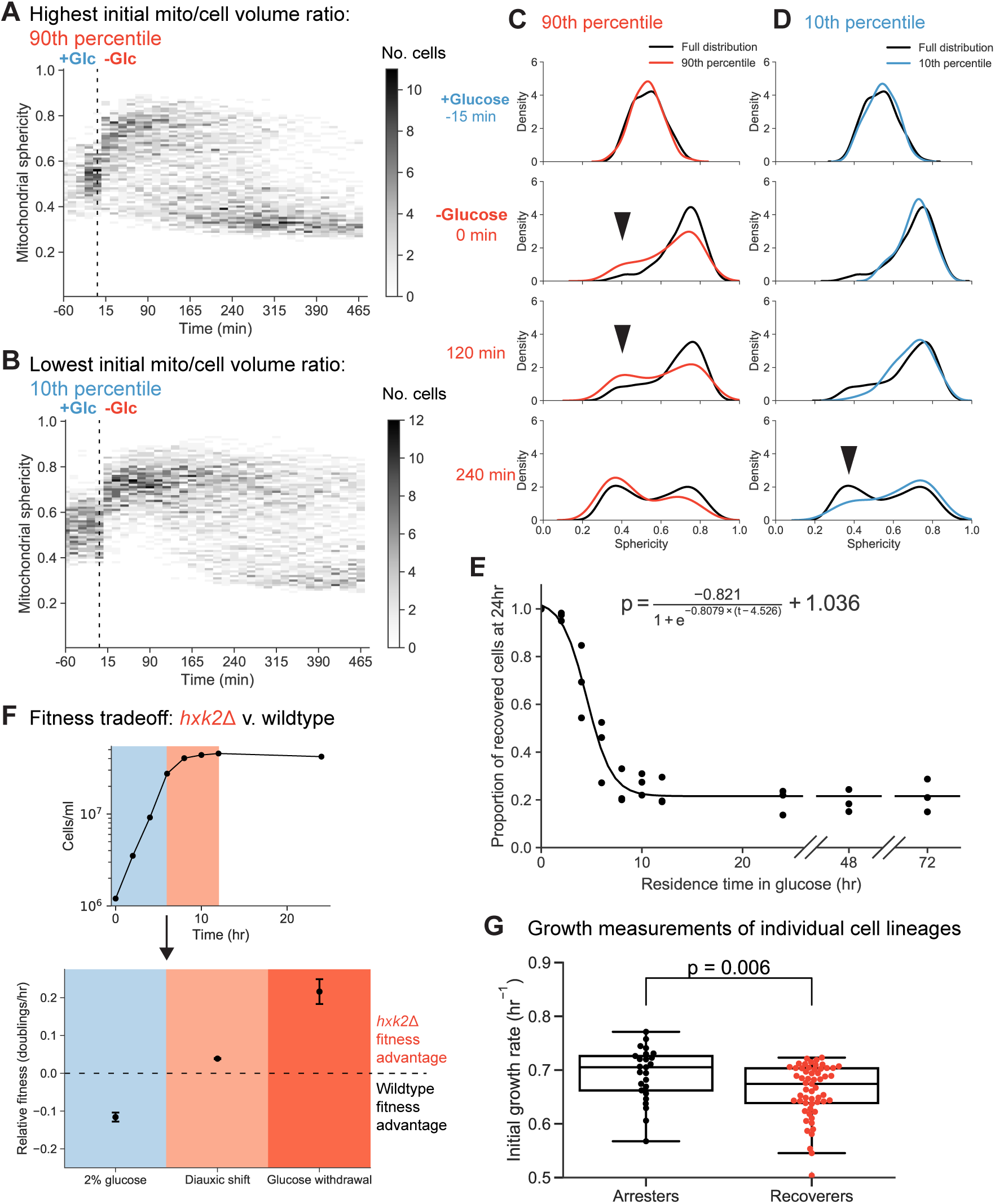
Post-starvation adaptation is associated with preparatory activity that imposes a short-term fitness cost. (A)-(B) Time-resolved heat maps of mitochondrial sphericity in the presence of high glucose and during acute glucose starvation for cells with the top 10% *(A)* and bottom 10% *(B)* mitochondrial to total cell volume ratios immediately prior to glucose washout (N = 133 and N = 132, respectively). Intensity reflects the absolute number of cells with a mitochondrial sphericity index within the given bin. (C)-(D) Histograms of selected timepoints from *(A)* and *(B)*, relative to the entire distribution. Arrows highlight discrepancies in relative subpopulation sizes. (E) Adaptation probability as a function of residence time in media containing high glucose prior to glucose disappearance. Cells expressing Hxt3p-mNeonGreen (yLB432) were grown in synthetic media lacking amino acids and containing non-fermentable potassium acetate as a carbon source, then switched into otherwise identical media containing high glucose for the specified number of hours prior to sudden glucose starvation. Recovery was scored as the presence or absence of Hxt3p-mNeonGreen signal by flow cytomery. The solid line represents the logistic function, displayed above, calculated from the data by non-linear least squares fit. Three biological replicates, N = 40,000 cells for each time point. (F) Upper panel: cell density of a culture initiated with equal proportions of wild-type (yLB365) and *hxk2*Δ (yLB373) in synthetic media containing high glucose. Blue and red shaded regions indicate exponential and diauxic growth phases, respectively. Lower panel: relative fitness of wild-type and *hxk2*Δ mutants during exponential phase growth and after the diauxic shift (as in upper panel) calculated as the relative rates of doublings/hr and measured by the changes in the ratios of the two genotypes by flow cytometry. Equivalent fitness measurement in the first 8 hours following sudden glucose deprivation is displayed in darker red. Error bars, one standard deviation. Three biological replicates, N = 40,000 cells analyzed by flow cytometry for all time points of all conditions. (G) Distribution of growth rates of single-lineage microcolonies in the presence of high glucose, partitioned by the failure or success of lineages to complete one doubling in the 12 hr following glucose starvation (arresters and recoverers, respectively). Mann-Whitney *U* test p-value = 0.006. See also Figure S5.

These data imply a hysteresis in metabolic flexibility. Cells with larger mitochondrial networks, weaker glucose repression, and some respiration adapt faster to sudden glucose deprivation. We wished to test this directly. For higher-throughput batch experiments, we identified the abundance of the hexose transporter Hxt3p as a marker of presence or absence of metabolic adaptation under our glucose deprivation assay. Hxt3p is highly expressed during exponential growth in fermentative conditions but is transcriptionally repressed and existing protein is transported to the vacuole and degraded during growth on nonfermentable carbon sugars (Perez et al., 2005; Roberts and Hudson, 2006; Snowdon et al., 2009). Recovering cells transport Hxt3p to the vacuole and degrade it, but in arresters, which fail to adapt, Hxt3p continues to reside on the plasma membrane. The behavior of Hxt3p after glucose starvation matched our expectations: it was completely lost from *hxk2*∆ cells, retained on all *snf1*∆ cells (save for phase-dark dying cells), lost on some wild-type cells that recovered and retained on those that did not (Figure S5A). We further validated the behavior of Hxt3p by flow cytometry and recapitulated the bimodal and unimodal distributions seen in our mitochondrial metrics (Figure S5B).

We asked how the fraction of recoverers within a population depended on the metabolic history of the culture. We induced cells expressing Hxt3p-mNeonGreen to respire by culturing them with a nonfermentable carbon source. We then transferred cells to media containing high concentrations of glucose for varying residence times in a state of fermentative growth before resuspending them in media lacking glucose. We performed these experiments in prototrophic strains and omitted amino acids from the both the glucose-containing and glucose-free media to prevent cell replication, allowing us to score the probability with which cells can successfully switch metabolic states based on the proportion of cells that have lost Hxt3p-mNeonGreen signal. We observed a decrease in adaptation capacity with increasing time in the presence of glucose, the dynamics of which appear to fit a logistic decay curve (r^2^ = 0.95) (Figure 5E). It is of note that, many generations after their ancestors last respired, the culture maintains a steady state at which roughly 20% of cells recover rapidly after an environmental shift.

If we posit that starvation recovery, or lack thereof, is a binary state in correspondence with the steady-state bimodality observed in both mitochondrial morphology and Hxt3p abundance, then all populations would consist of some combination of the two cell types. A culture grown from a small number of founding cells on a nonfermentable carbon source is by definition composed solely of the recoverer type, as this metabolic state is a prerequisite for growth and division in this context. These cells have already successfully assumed a respiratory metabolic state. As we observe a decrease in recovery probability with increasing exposure time to glucose, we thus see interconversion between the two states as a previously homogeneous population develops heterogeneity, and the decrease in recovery probability reflects the conversion rate between recoverer and arrester metabolic states. By fitting our data to a logistic curve, we estimate this rate to be 0.23 hr^−1^.

A culture maintained in exponential phase in glucose-rich media for 24 hr or more has reached an asymptote on the metabolic memory curve. We expect that this probability of recovery reflects the steady state ratio of arrester cells to recoverer cells in the absence of environmental cues, when switching in both directions (from recoverer to arrester and vice versa) occurs at equal rates. We calculate that this ratio is 3.65, allowing us to infer that the rate at which arrester cells prepare for respiration and switch metabolic states must be 0.23 hr^−1^ / 3.65 = 0.06 hr^−1^. We measured these rates directly by growing individual founder cells into extended lineages in our microfluidics device and detecting switching events through inconsistencies in cell pedigrees once these lineages were subjected to sudden starvation (see Methods). We calculated the recoverer-to-arrester switching rate to be 0.18 ± 0.02 hr^−1^ and the arrester-to-recoverer switching rate as 0.08 ± 0.02 hr^−1^, both consistent with the values derived from our bulk assays.

Why do all cells not retain the metabolic flexibility to adapt to sudden environmental perturbations, given the obvious advantage of recovery? Our data suggest that recoverer cells are prepared for starvation due to partial glucose derepression, residual levels of machinery necessary for growth in derepressing environments, outright respiratory activity, or some intermediary status on the path to fully respiratory metabolism. If so, in permissive environments featuring abundant glucose, such prepared cells would generate ATP more slowly or bear protein or transcriptional loads that other cells would not. Recoverer cells would grow and replicate more slowly to maintain their metabolic flexibility. As long as the glucose concentration remains high, they would sacrifice current growth potential for resistance to a future, hypothetical period of glucose starvation. Previous work has identified such a fitness tradeoff during shifts between two carbon sugars, glucose and maltose, in *hxk2*Δ mutants (New et al., 2014).

We tested for context-dependent fitness differences between wild-type cells and uniformly-recovering *hxk2*Δ mutants. We inoculated cultures with equal numbers of cells of each genotype, each bearing a different fluorescent label. We tracked their relative changes in abundance in synthetic media containing glucose during exponential phase growth and through glucose exhaustion at the diauxic shift. In separate cultures, we performed glucose withdrawal and measured the relative effect on each strain.

We found that *hxk2*Δ cells display a fitness defect of approximately 12% during growth in glucose-containing media but have a fitness advantage, relative to wild-type cells, of 3.9% after rapid exponential growth ceases (Figures 5F, S5C, and S5D). During sudden starvation, *hxk2*Δ cells outgrow wild-type cells by 21.6%. These reciprocal relative fitnesses in the presence or absence of glucose are consistent with our hypothesis.

We next examined whether a reciprocal fitness effect is detectable within wild-type populations. Single-cell growth rates are difficult to measure in *S. cerevisiae* due to the nature of budding, in which much of the earliest growth is limited to tiny, difficult-to-resolve daughter compartments. We instead pooled measurements across microlineages, tracking the total area encompassed by individual founding cells and their progeny across several cell divisions and assessing the collective growth of the pedigree before and following sudden glucose washout. This method is sensitive to the ability of cells to switch metabolic states at any point within the lineage, which, though rare, presents a confounding effect. We were, however, able to distinguish clearly between groups of cells in which all members collectively either succeeded or failed to complete one round of division following 12 hr of starvation (Figure S5E). We partitioned these microcolonies into two states, recoverers and arresters, and compared the pre-starvation growth rates across the two groups. The recoverer and arrester groups were enriched in cells that were slower- and faster-growing, respectively, in the original, high-glucose environment (p = 0.006, Figure 5G). These data indicate that cells best able to adapt during sudden glucose deprivation grow more slowly in high glucose.

### Bimodal starvation behavior with natural variation in steady states in wild yeast isolates

If cells must balance their short-term competitive fitness against long-term stress resistance, then we might expect that strains dwelling in distinct niches would all obey a similar fitness tradeoff but would evolve to establish distinct distributions of metabolic types in response to their particular environmental challenges--there should exist ecological diversity in extent of risk or risk aversion which should match strains’ metabolic set points. To test this, we integrated our Hxt3p-mNeonGreen marker for metabolic adaptation into a series of ten additional *Saccharomyces cerevisiae* strains isolated from a range of contexts (Liti et al., 2009). We measured their pairwise fitnesses in the presence of high glucose and following sudden glucose starvation, using one strain as a common reference competitor.

We observed an inverse correlation between their relative fitness in the presence or sudden absence of glucose across all strains (slope = −0.71, r^2^ = 0.84) (Figure 6A). We further examined our fluorescent markers, for both mitochondria and Hxt3p, the diagnostic hexose transporter, in a natural yeast isolate during microfluidics-based glucose withdrawal, observing similar bimodality underlying arrest and recovery behavior during sudden starvation (Figures 6B and 6C). We conclude that the presence of two physiological states, recoverers and arresters, which either maximize their preparation for sudden changes in carbon sources or their rate of proliferation on glucose, extends beyond laboratory cultivation and that the balance between them is likely to have been shaped by local ecological forces.

**Figure 6.**
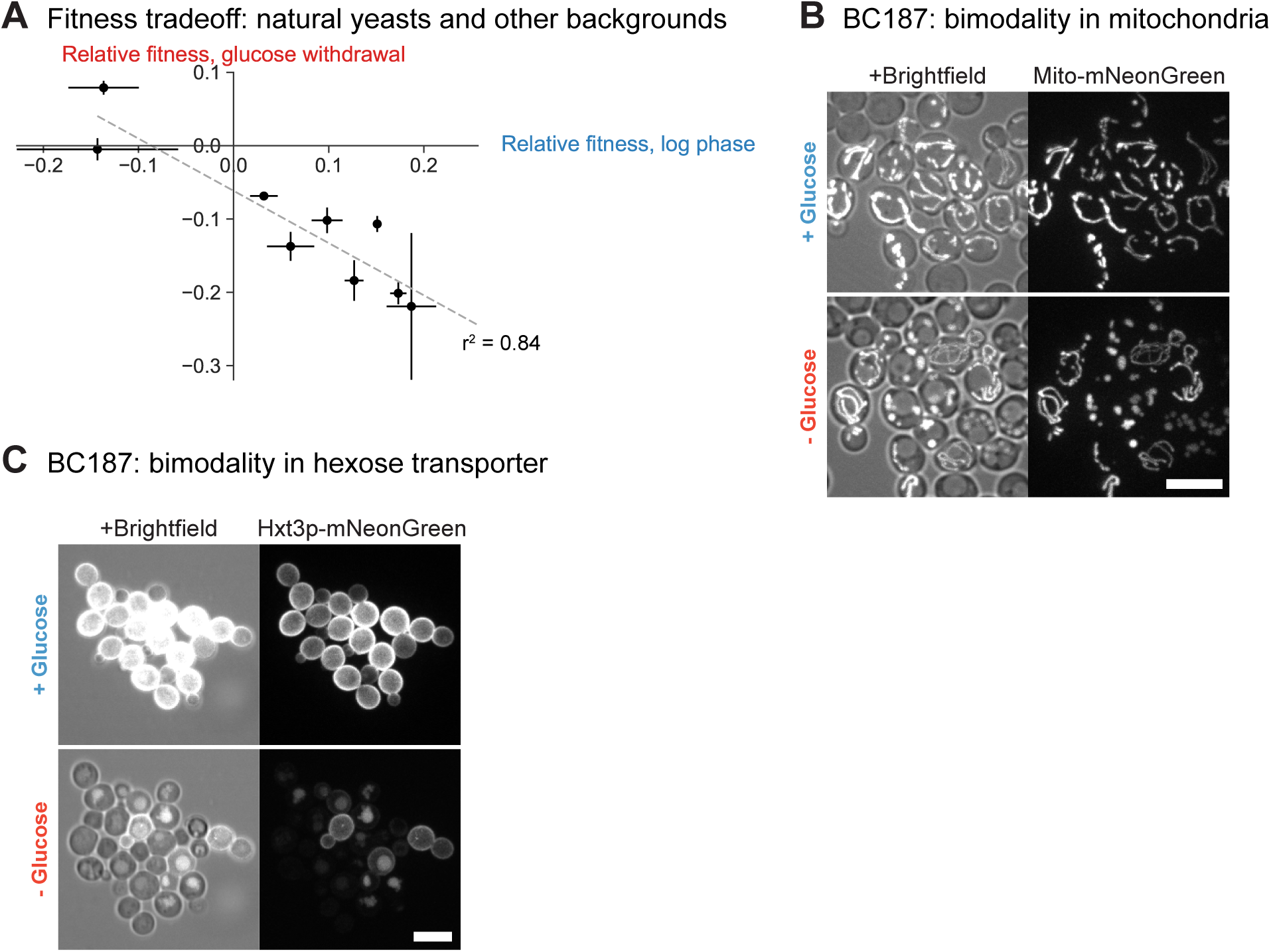
Natural variation in starvation behavior obeys a fitness tradeoff. (A) Relative fitnesses of diploid derivatives of YJM978 (yLB463), CEN.PK (yLB467), DBVPG1373 (yLB470), L-1374 (yLB474), BC187 (yLB478), Y12 (yLB486), K11 (yLB492), YPS606 (yLB494), and UWOPS83-787.3 (yLB496) expressing mNeptune under the *ACT1* promoter, measured against a common reference, YS2 (yLB480), when assayed in pairwise competitions, both in exponential phase growth in the presence of abundant glucose and following abrupt glucose deprivation. Three independent biological replicates measured for all competitions. Error bars, one standard deviation. (B)-(C) Microscopic images of mitochondrial matrix marker *(B)* and Hxt3p-mNeonGreen *(C)* in diploid BC187 derivative (yLB453), in synthetic media containing high glucose and 7 hr following abrupt glucose withdrawal. Scale bars, 10 μm.

## DISCUSSION

We investigated the response of budding yeast cells to sudden glucose starvation using mitochondrial abundance and morphology as proxies for a cell’s metabolic state. Glucose starvation causes mitochondrial collapse in the majority of cells in conjunction with a fall in intracellular pH and electric potential across the mitochondria. As starvation continues, cells respond bimodally, reflecting an inherited state: recoverers return to a normal mitochondrial morphology, increase their mitochondrial content, raise intracellular pH and mitochondrial potential, and resume growth, while arresters remain arrested with collapsed mitochondria. Mutations that affect metabolic regulators can lock cells in either state, and recoverers grow more slowly in the presence of high glucose but have a strong fitness advantage during the switch from glucose to non-fermentable carbon sources. These two states are present in wild yeast strains and variation in their relative proportions suggest that the presence of two phenotypes in a population of genetically identical cells is a form of bet-hedging.

Our mitochondrial morphological metrics are easily quantifiable properties that are disrupted in tandem with a host of additional cellular structures, making them valuable reporters on global organizational status. Our sphericity index captures intracellular changes on multiple levels: it reports on the state of shock associated with the majority of cells in the early stages of starvation, it declines in cells that recover and resume growth producing a bimodal distribution that separates recovering cells from those that have not started recovery, and it reports on the increased tubularity and branching of the mitochondrial network in cells growing on suboptimal carbon sources. The gradual increase in mitochondrial volume ratio in adapting cells also allows us to track recovery as cells reorganize their internal architecture to exploit a different carbon source using different metabolic pathways. Crucially, both of these parameters signal imminent changes in metabolic state well before any detectable increases in total cell volume or bud production, which lag behind mitochondrial remodeling by more than one hour.

Based on the correlations between cell growth rates with mitochondrial morphology, intracellular pH, and mitochondrial potential, we propose a model of budding yeast metabolism that we illustrate in Figure 7. We argue that genetically homogeneous populations growing in the presence of abundant glucose are bimodal in rates of glycolysis and in broader metabolic state, with one subset of cells, arresters, fermenting heavily and another subpopulation, recoverers, displaying partial glucose derepression and respiration. When glucose is suddenly removed, arresters fail to adapt their metabolism to the new nutrient limitation and cannot grow. Glucose starvation produces stress, as manifested in the dramatic and uniform decrease in cytosolic pH in both recoverers and arresters. In arresters, which were producing all their ATP from glycolysis, glucose-starvation produces a profound drop in ATP production which cannot easily be remedied, leading to a prolonged arrest without functional mitochondria. In recoverers, however, respiration is already occurring, allowing them to respire amino acids in the medium (and possibly any reserve carbohydrates that they have accumulated) to restore intracellular ATP levels and resume growth.

**Figure 7.**
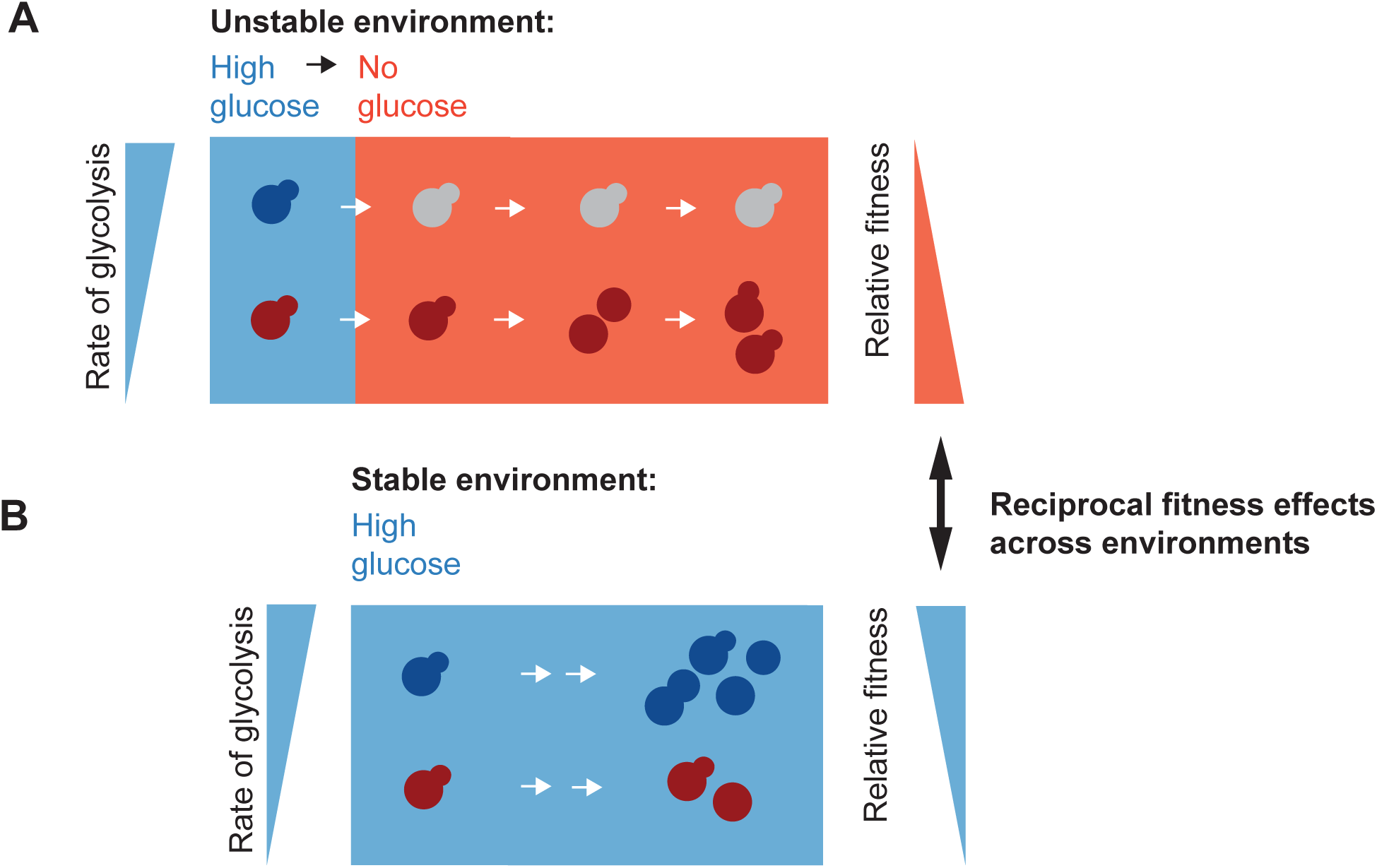
A model of mitochondria and metabolic transitions. (A) A clonal population of yeast growing in the presence of abundant glucose (blue background) is composed of cells in two distinct metabolic states in which the relative rate of glycolysis is either high or low (blue and red cells, respectively). Upon sudden glucose starvation (red background), cells with high glycolytic activity arrest indefinitely, while those with low activity adapt and ultimately resume growth. Lower initial rates of glycolysis result in a post-starvation fitness advantage. (B) In stable, high-glucose conditions, maximizing the rate of glycolysis (blue cells) can sustain a higher growth rate and thus produce a fitness advantage.

Although recoverers have an advantage upon glucose starvation, arresters grow faster at high glucose concentrations, most likely from a combination of investing less in producing mitochondria and a higher rate of ATP production from glycolysis. The two metabolic strategies differ in their relative efficacy in different environments: recoverers are prepared for a future environmental shock at the cost of a lower growth rate on glucose, while arresters have a severe fitness disadvantage following sudden glucose starvation (Figure 5G). One metabolic strategy emphasizes carbon source specialization, utilizing high levels of glucose as efficiently as possible while they last. The other strategy is generalist, predicated on growing more conservatively in rich environments to retain growth potential during environmental instability. That an isogenic population contains cells of each type, those which are more competitive in the current environment and those which will be more resistant to unexpected stresses, helps to maximize the population’s fitness over a wide range of future environments.

What determines the state of an individual cell? We have demonstrated that daughter cells are likely to display the same phenotype as their mothers, indicating that metabolic state is heritable. Our collective experiments have ruled out, in addition to superstable metabolic memory of previous metabolic cues, age dependence (given the heritability of our phenotype, as yeast daughters are born young at the expense of their mothers), cell cycle status, and cell size in the determination of these states. We observe these state ratios in well-mixed liquid cultures as well as in microfluidic chambers, precluding spatial effects in cell state establishment. We therefore propose that metabolic state switching occurs stochastically. The simplest model is that rates of switching between the two states are constant and independent of the environment. When cells are glucose-starved, only recoverers can grow, leading to a population made almost entirely of recoverers. When this population is shifted to glucose, recoverers transition into arresters until the population reaches a steady state in which the rates of switching between the two states are equal. It is possible that the rate of switching from recoverers to arresters is reduced in respiring cells, but the difference in the growth rate of the two populations makes measuring this rate difficult.

The adaptive value of recoverer cells, “persisters” which have prepared for starvation in by reducing their current growth rate, depends on the probability of a sudden environmental challenge. If no such event arrives, recoverers reduce the population’s overall fitness. Evolutionary forces should shape the switching rates between the two states as a function of the relative frequency with which a population with access to abundant glucose experiences sudden starvation: more frequent starvation should be associated with a higher steady state frequency of recoverers. We have observed population-level correlations between relative fitness advantages and disadvantages across rich and starvation conditions for a collection of yeast isolates, indicating that these strains all obey a similar fitness tradeoff between recoverers and arresters but indicating that there is natural variation in the ratio of carbon utilization strategies within a similar bimodal framework.

Our work reveals heritable, non-genetic individuality within yeast populations that balances current fitness concerns against future environmental uncertainty. Further, while previous studies have observed natural, inter-strain variation in the response to environments containing multiple carbon sugars (Escalante-Chong et al., 2015; Lee et al., 2017; Roop et al., 2016; Wang et al., 2015), our observations extend the variation into a non-genetic context in which starvation is not preceded by any predictive cues.

Is the behavior we have observed bet-hedging? Though the term is often applied colloquially, it was introduced to refer to the behavior of an isogenic population that has evolved in unpredictably fluctuating environments to maximize long-term fitness and reduce variability in the number of cells or progeny produced over time (Gillespie, 1974; Slatkin, 1974). This hedging can involve a single, conservative phenotype that is suboptimal in multiple environments but maladaptive in none, or diversification into multiple phenotypes that are individually advantageous in some environments and detrimental in others but collectively mitigate risk across all environments (Philippi and Seger, 1989; Seger, 1987). Crucial to this definition of bet-hedging is its origins in the selective pressures applied by an unstable environment lacking predictive cues. In such environments, organisms should optimize the geometric mean fitness over a long period of fluctuation, weighted for the magnitude and frequency with which organisms experience each environment. The inability to measure the underlying environmental dynamics as experienced by such organisms over time makes bet-hedging difficult to demonstrate empirically (de Jong et al., 2011).

Our experiments identify phenotypic heterogeneity within clonal populations that produces two cell states, each of which displays a fitness advantage in one of two environments. In addition, strains from the wild show obey similar trade-offs between growth in abundant glucose compared to growth after glucose starvation, suggesting that this trade-off is under natural selection, As it stands, this makes our study of heterogeneity in the response to glucose starvation the best evidence to date for the existence of bet-hedging in budding yeast (see the history of bet-hedging claims and an associated ranking system in Simons, 2011).

When the dynamics of natural environmental fluctuations are unknown, as is the case for our natural yeast strains, definitive proof of bet-hedging behavior is impossible. However, our nutrient-withdrawal paradigm, applied in the context of experimental evolution, would permit direct testing of both the existence of bet-hedging and, if applicable, the genetic mechanisms by which the frequencies and properties of recoverers and arresters are controlled. By evolving yeast under conditions that fluctuate between glucose-rich media and sudden glucose starvation, with nutrient shifts occurring at varying average frequencies, it would be possible to determine whether the rate of metabolic state switching between cell types optimizes the geometric mean fitness determined by the rate of environmental instability. If successful, this could offer insight into the mechanisms that control this bimodality, the rates of interconversion between states, and the nature of bet-hedging in budding yeast.

## ACKNOWLEDGMENTS

This work was supported by NIH grants DP2AI117923-01 to E.C.G., R01GM043987 to A.W.M. and the NSF/Simons Center for Mathematical & Statistical Analysis of Biology at Harvard (#1764269 (NSF) and #594596 (Simons Foundation)). L.E.B. was supported by the National Science Foundation Graduate Research Fellowship under Grant No. DGE1144152. The authors thank the Harvard Bauer Core Facility for technical assistance. We thank P. Stoddard, S. Srikant, Y. Sun, and other members of the Murray and Garner labs for helpful discussions. The authors declare no competing interests.

## AUTHOR CONTRIBUTIONS

Conceptualization, L.E.B., Q.A.J., E.C.G., and A.W.M.; Methodology, L.E.B., E.C.G., and A.W.M.; Investigation, L.E.B.; Writing and editing, L.E.B., E.C.G., and A.W.M.; Supervision, E.C.G. and A.W.M.

