## Supplementary materials for "Bet hedging buffers budding yeast against environmental instability"

**This supplement includes:**

Materials and methods
Figures S1-S5
Tables S1-S2
Legends for movies S1-S3
References for materials and methods

**Other supplementary material for this manuscript includes the following:**

Movies S1-S3

### MATERIALS AND METHODS

#### Data and code availability

Mitochondrial morphological data and code used in the analysis and production of figures are available at https://github.com/bagitmery.

#### Yeast strains and media

All experiments were performed using strains constructed in a modified W303 background in which the *bud4* and *rad5-535* alleles were replaced with their respective, functional loci from the S288C background (*MATa leu2-3,112 rp1-1 can1-100 ade2-1 his3-11,15 BUD4-S288C RAD5*). The mito-mNeonGreen construct was derived from pVT100U-mtGFP, a gift from Benedikt Westermann (Addgene plasmid 45054)

(Westermann and Neupert, 2000) and consists of a pRS403 integrating vector (Sikorski and Hieter, 1989) harboring an *ADH1* promoter, the first 69 amino acids of subunit 9 of the *Neurospora crassa* F0 ATPase (the mitochondrial presequence preSu9), a short linker sequence (Sheff and Thorn, 2004), mNeonGreen (Shaner et al., 2013), and an *ADH1* terminator. All sequences were codon-optimized for expression in yeast.

Integrating plasmids carrying fluorescent pH reporters were constructed by introducing a yeast-optimized sequence encoding ratiometric pHluorin2 (Mahon, 2011) and an *ADH1* terminator into the pRS403 integrating vector. In the mitochondrial pH reporter, pHluorin2 was placed under the control of the *ADH1* promoter, directly downstream of preSu9; in the cytosolic pH reporter, expression was controlled by the *ACT1* promoter.

All other fluorescent fusion constructs consist of a common linker and a fluorescent protein joined to the C terminus of the labeled protein, expressed from its native locus and under its native promoter. Fusion sequences were joined by isothermal assembly (Gibson et al., 2009). All constructs and markers for both fluorescent labeling and gene disruptions were introduced by standard yeast genetic methods (Amberg and Strathern, 2005) and confirmed by PCR and, where applicable, DNA sequencing.

Respiratory-deficient petite strains were produced by incubating cells in 25 μg/ml ethidium bromide for 24 hr. Loss of mitochondrial DNA was verified by DAPI staining and the absence of growth on non-fermentable carbon sources.

Experiments were performed in synthetic complete (SC) media prepared from 10x yeast nitrogen base (YNB); 100x stocks of adenine, tryptophan, and uracil; 2 g/L complete amino acids lacking the three amino acids above; 10x stock of the specified carbon source, if any; and sterilized water. For bulk assays, YNB was prepared according to the Wickerham specifications (Wickerham, 1951); for microscopy assays, this recipe was modified to omit riboflavin and folic acid, the two primary contributors to autofluorescence (Sheff and Thorn, 2004). Where noted, select bulk assays and microfluidics-based rate-switching assays were performed in minimal media lacking all amino acids but otherwise prepared as described for SC media. Ingredients for the preparation of these media were purchased from Sigma-Aldrich.

For growth and starvation behavior assays, strains were streaked from stocks stored in 15% glycerol and sterilized water at -70°C onto agar plates of the appropriate selective dropout media. After two days of growth at 30°C, single colonies were used to inoculate cultures that were continuously maintained in early exponential phase (a density of less than $5 \times 10^6$ cells/ml as measured on a Beckman Coulter Counter), also at 30°C, for a minimum of 24 hr and no more than 36 hr (unless otherwise noted), prior to the initiation of any experiment. All experiments consist of a minimum of three biological replicates performed on separate days.

**Imaging sample preparation and microfluidics**

Single-timepoint images of cells grown in varying carbon sources were imaged in liquid media in 96-well plates with optical glass bottoms (Grenier Bio-One), precoated with 0.5 mg/ml concanavalin A (MP Biomedicals) in sterile water.

Cells examined during and following abrupt nutrient shifts were immobilized in silicone CellASIC Y04 plates for haploid yeast cells, and the flow of media was controlled with the CellASIC ONIX microfluidic system (EMD Millipore). Plates were pretreated through perfusion of 0.5 mg/ml concanavalin A, freshly dissolved in a buffer of 10 mM $Na_2HPO_4$ and 0.5 mM $CaCl_2$, pH 6.5, immediately prior to use. Priming with this solution was performed at an initial flow rate of 5 psi for 5 min, followed by a 90 min perfusion at 2 psi, then washout with yeast media for 5 min at 5 psi. Cell cultures, prepared as described above, were loaded into the chamber at 4 psi for 10 sec. Cells were continually supplied with growth or starvation media at a rate of 2 psi thereafter; based on the manufacturer's specified flow properties, we estimate this to result in roughly 4.5 complete media exchanges per minute within the cell chamber. Cells were allowed to equilibrate within the chamber for a minimum of 1 hr before the primary carbon source was removed.

For measuring mitochondrial potential, cells were incubated with 0.5 μM MitoTracker Red CM-$H_2$Xros in the dark for 15 min prior to imaging or flow cytometry.

**Microscopy**

Images were acquired using a Nikon TI inverted microscope equipped with a Yokogawa CSU-10 dual spinning disk confocal unit, 16.0 μm-pixel Hamamatsu ImagEM EM-CCD camera, Nikon 100x NA 1.45 TIRF objective, and MetaMorph software. 488 and 594 nm lasers were used with 525/45 nm (GFP) and 609/57 nm (RFP) bandpass filters for imaging of green (mNeonGreen) and red (mNeptune) fluorophores, respectively, with exposure times of 122.12 ms. Z-stacks were collected in 0.2 μm slices with a minimum of 8 μm total depth. The two-dimensional images shown were flattened by maximum intensity z-projection. Microscopy was performed at room temperature.

**Image analysis**

Brightfield images of cells were segmented with the CellStar algorithm run via MATLAB plugin (Versari et al., 2017). The resulting cell masks were manually inspected and corrected in the event of over- or under-segmentation. Cell volume was calculated by fitting the generated cell contour to an ellipse and projecting into three dimensions based on the geometry of a prolate ellipsoid. Buds smaller than 20 $\mu m^3$ were excluded from analysis; this was determined to be the threshold for the generation of reliable masks by the manual examination for > 200 cells. Tracking, also performed by the CellStar algorithm, was used to assign unique identification numbers to all masks and allow for analysis of single-cell trajectories. Prior to mitochondrial analysis, cell masks were used to crop images and generate single-cell z-stacks. Background noise as calculated within a five-pixel band about the cell contour was used to populate all pixels outside of the cell mask. Cell volumetric calculations and preprocessing for mitochondrial analysis were performed by custom MATLAB scripts available at https://github.com/bagitmery.

Mitochondrial segmentation was performed using MitoGraph software (Viana et al., 2015). Three-dimensional reconstructions of mitochondrial content in Visualization ToolKit (VTK) format were imported into a Python 2.7 environment via the Python VTK wrapper library. Mitochondrial and whole-cell volumetric data were matched by cell identification numbers and statistical analysis and plotting were performed in Python. Mitochondrial/cell volume ratios were calculated as the total mitochondrial volume divided by the total cell volume. For sphericity index, the volume of each mitochondrion was used to calculate the surface area of a sphere of equivalent volume, which was divided by the actual surface area of that mitochondrion. This value was calculated for all topologically distinct mitochondrial units individually, and an overall sphericity index was calculated as the average of all distinct mitochondrial sphericity values within a cell, weighted by the volume of each mitochondrion. Individual mitochondrial units of less than 45 $nm^3$ total volume were flagged as potential artifacts; this was determined to be the threshold of reliable segmentation by the manual examination of > 200 cells. Cells containing five or more such fragments were excluded from analysis.

Cells' budded or unbudded state, mother-daughter relationships, and time of visible growth resumption during glucose starvation were annotated manually and integrated with mitochondrial data and analyzed in Python.

Composite images, maximum intensity z-projections, and movies were assembled for presentation using the Fiji distribution of ImageJ (Schindelin et al., 2012).

**Bulk glucose withdrawal**

Cells grown in batch culture conditions were starved of glucose after allowing a clonal culture, derived from a single, freshly streaked colony, to grow exponentially for a minimum of 24 hr in the presence of high glucose as described above. Cells in 5-10 ml culture volumes were collected by centrifugation at 600 rpm for 5 min, washed twice with 50 ml media lacking glucose, and resuspended in 5-10 ml of glucose-free media. At regular intervals, samples were analyzed by flow cytometry as described below.

**Flow cytometry**

Samples were prepared for flow cytometry by treating 1 ml aliquots of cell culture with cycloheximide (Sigma-Aldrich) added to a final concentration of 100 µg/ml. Samples were stored at 4°C in the dark until analysis, performed within one day. Cycloheximide stocks were prepared at 100x in ethanol and stored at -70°C for no more than one year. Analysis was performed on an LSR II flow cytometer (BD Biosciences) using the 488 nm laser and a 505 nm long pass filter and 530/30 nm bandpass filter to measure mNeonGreen intensity. For pH assays, samples were analyzed live, in the original media environment and without additional treatment, using the 405 nm and 488 nm lasers, each coupled with a 505 nm long pass and 530/30 nm bandpass filter. For all experiments, 40,000 cells were analyzed. Data were imported into the Python 2.7 environment using the FlowCytometryTools package and custom Python scripts were used to isolate the single-cell population on the basis of forward and side scatter profiles and to examine the distributions of fluorescent signals.

**Calibration and measurement of pH**

*In situ* calibration of the pHluorin2 ratiometric reporter was performed by collecting cells expressing the pHluorin2 construct and growing them in exponential phase in synthetic complete medium, resuspending them in PBS, and treating with 100 µl/ml digitonin, to permeabilize the plasma membrane, for 10 min. Cells were then separated into nine aliquots and resuspended in a series of citric acid-$Na_2HPO_4$ buffers spanning pH 5.0-9.0 in half-pH-unit intervals. Cells were analyzed immediately by flow cytometry as described above. The relative signals obtained from the 405 nm and 488 nm laser settings in three biological replicates were plotted against pH and fit to an exponential function. In subsequent live-cell experiments, this standard curve was used to compute pH from relative 405 nm / 488 nm signal intensity.

**Oxygen consumption measurements**

Oxygen depletion was measured in 96-well microplates containing embedded oxygen optical sensors (OxoPlate OP96C, PreSens). Cells were grown for 24-48 hr in log phase in synthetic media containing 2% glucose (or other carbon sugars as noted) and added to microplate wells at a density of $4 \times 10^6$ cells/ml. Sensor output was measured using a Synergy Neo2 plate reader (BioTek) with a monochromator set for excitation at 540 ± 2 nm and emission at 509/20 nm (reference signal) and 650/20 nm (indicator signal). Fluorescent signal and absorption at 600 nm were measured every 15 min for 8 hr at 30°C with plate shaking between time points. Oxygen saturation was computed from the ratiometric signal according to the manufacturer's instructions, using air-saturated and oxygen-depleted water (0.01 g/ml $Na_2SO_3$) as calibration standards, measured in parallel with the biological samples. Oxygen depletion rates were calculated by normalizing oxygen readings to cell density, performing a linear regression on the linear portion of the depletion trace by least squares fitting, and reporting the obtained slope. Slopes were measured for three biological replicates, each consisting of four technical replicates.

**Starvation phenotype state-switching measurements**

The rates of interconversion between starvation phenotypes were calculated from microfluidics experiments in which cells expressing Hxt3p-mNeonGreen (yLB432) were pregrown in synthetic medium lacking amino acids and containing 2% glucose for 6 hr, after which the flow was replaced with identical medium containing 0% glucose. Cells were classified as arrested or recovering on the basis of their retention or turnover, respectively, of Hxt3p-mNeonGreen 6 hr post-glucose withdrawal. The birth state of each cell was inferred by comparing its phenotype to those of its ancestors and descendants and assuming the fewest state switches necessary to produce the observed phenotypic patterns within a lineage. These data, along observation time between each cell's birth and its sudden starvation, were collected for a total of 2,702 cells across five independent experiments. These data were partitioned based on the initial phenotype of each cell at birth, and the proportion of cells not experiencing a phenotypic switch was calculated for all cells with a shared observed lifetime (binned into 15-min increments, the frame rate at which fluorescent images were collected). These probabilities were fit to the k = 0 case of the Poisson distribution in which the average number of events per interval is a product of the switching rate and the length of the observation window. We calculated a recoverer-to-arrester switching rate of 0.18 $\pm$ 0.02 $hr^{-1}$, with our data fitting a Poisson probability mass function with $r^2 = 0.77$. For arrester-to-recoverer switching, we restricted our analysis to cells with lifetimes of 90 min or less, as the probability of switching decreased after 90 min, likely due to the relatively high probability of cells switching back to the arrester phenotype during longer observation times. We calculated an arrester-to-recoverer switching rate of 0.08 $\pm$ 0.02 $hr^{-1}$, with $r^2 = 0.70$.

**Single-lineage growth measurements**

The MATLAB implementation of the Canny edge detection algorithm was used to determine the boundaries of microcolonies within brightfield microscopic images. Edges were subsequently connected by dilation. The two-dimensional area circumscribed by these boundaries was measured during growth in the presence of high glucose and following glucose deprivation, performed by microfluidics as described above. Pre-starvation growth rate was calculated by performing a linear regression on the increase in base two logarithm of observed area over time in the presence of high glucose.

**Competition assays**

Strains labeled with either mNeonGreen of mNeptune, expressed under the *ACT1* promoter from a construct integrated at the *HIS3* locus, were grown separately in log phase in complete synthetic media containing 2% glucose for a minimum of 24 hr prior to fitness testing. Cells were mixed at a 1:1 ratio in similar complete synthetic media at a density of 4 x $10^6$ cells/ml and either maintained in the presence of high glucose or subjected to bulk glucose withdrawal as described above. Samples were extracted every 1-2 hr, treated with 100 µg/ml cycloheximide and analyzed promptly by flow cytometry, with the ratio of each cell type calculated from clustering based on a Gaussian mixture model with full covariance matrix, implemented via the scikit-learn library in Python 2.7. Relative growth dynamics were quantified as the ratio of doublings per hour.

### SUPPLEMENTARY FIGURES AND LEGENDS

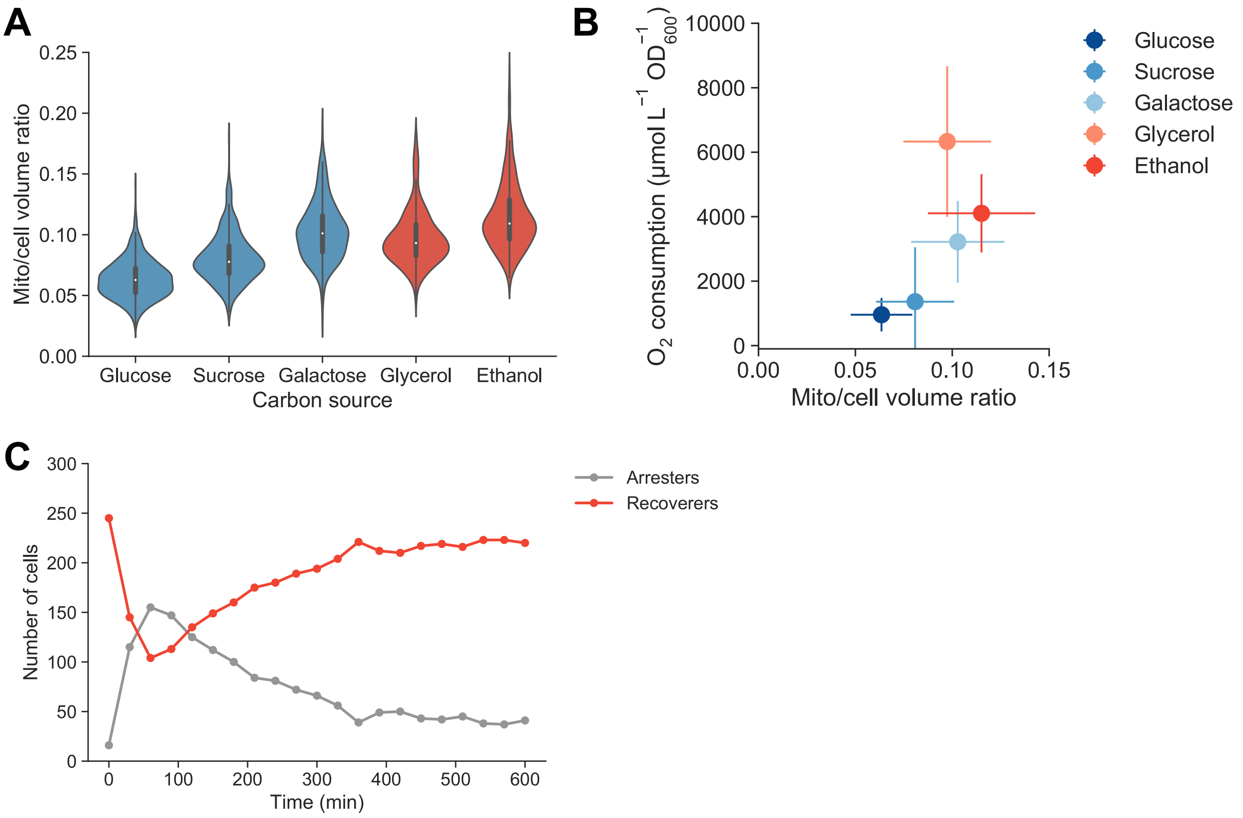


**Figure S1. Mitochondrial structure and metabolic output vary across environments. Related to Figure 1**

(A) The ratio of mitochondrial volume to total volume of cells expressing mitochondrially targeted mNeonGreen (yLB126) and growing exponentially in synthetic media containing the specified carbon sources, examined by microscopy. N ≥ 316 cells collected across three biological replicates. Fermentable carbon sources are shown in blue, non-fermentable ones in red.

(B) Oxygen consumption rates of yLB1 were measured in sealed cell culture plates equipped with a fluorescent oxygen sensor while cells grew exponentially in synthetic media containing varying carbon sources, conducted in parallel with volumetric measurements collected by microscopy in *(A)*. Rates normalized by doubling time of each strain, measured in tandem. Error bars indicate one standard deviation.

(C) Cells were classified as arrested or recovered on the basis of mitochondrial network morphology (sphericity index > 0.7 or ≤ 0.7, respectively) for 10 hr following glucose withdrawal. N = 261 cells collected across three independent biological replicates. A minority of cells displayed no mitochondrial collapse on the time intervals at which images were acquired (15 min). We are uncertain whether these cells experienced a temporary mitochondrial shock which is not detectable at this temporal resolution, and thus in our downstream analyses we simply classify cells as either recovering or arresting on the basis of their ultimate fate.

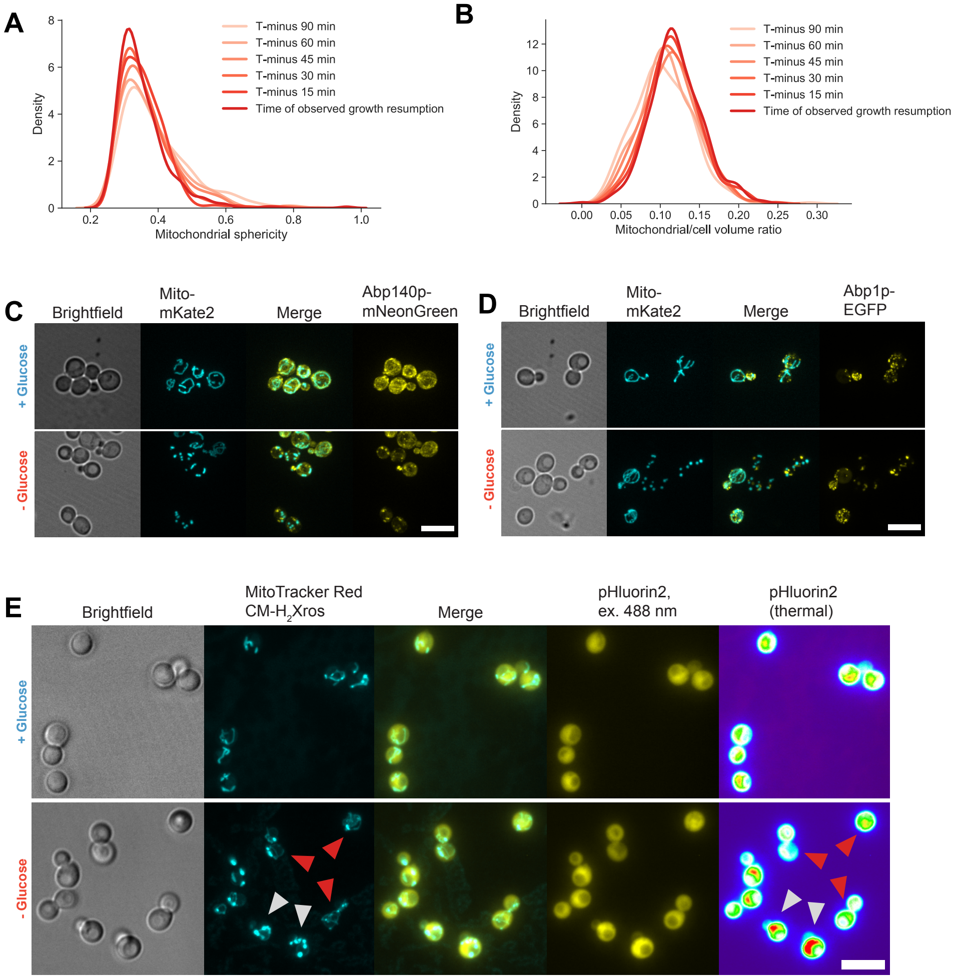


**Figure S2. Structural defects associated with growth arrest during sudden starvation. Related to Figure 2**

(A)-(B) Dynamics of mitochondrial sphericity *(A)* and mitochondrial/cell volume ratio *(B)* in N = 357 cells in the 90 min prior to visible growth resumption.

(C)-(D) Representative images of cells co-expressing a mitochondrial marker in tandem with actin cable-associated protein Abp140p (yLB69) *(C)*, or endocytic actin patch marker Abp1p (yLB45) *(D)*, growing in synthetic media containing high glucose and following glucose washout. Scale bars, 10 µm. The two actin-binding proteins largely maintain their distribution in cells where mitochondria are tubular and lose it in cells whose mitochondria collapse.

(E) Images of cells expressing ratiometric pHluorin2 (yLB397) and stained with MitoTracker Red CM-$H_2$Xros, in synthetic media containing high glucose and post-glucose deprivation. MitoTracker Red images are presented as maximum-intensity z-projections for clarity of mitochondrial morphology; pHluorin2 images consist of the summed intensity across all z-slices for retention of expression information. Excitation of pHluorin2 at 488 nm produces a stronger fluorescence signal as pH decreases. Far right panels depict pHluorin2 signal as a thermal heat map for ease of comparison. Red and gray arrows indicate examples of cells with tubular and collapsed mitochondria, respectively. Scale bar, 10 μm.

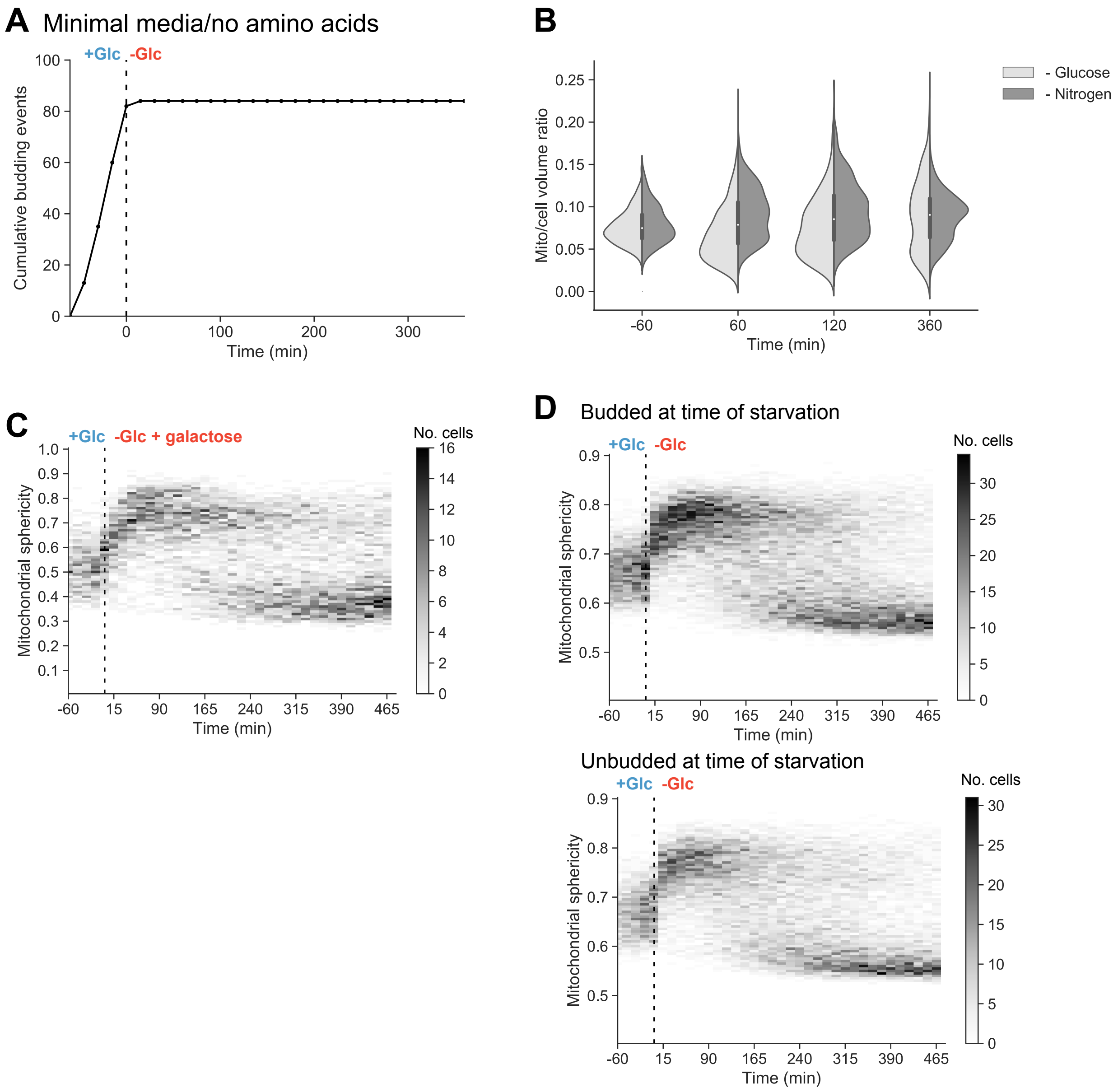


**Figure S3. Mitochondrial starvation heterogeneity is unique to glucose deprivation and is not explained by cell cycle starvation heterogeneity. Related to Figure 3**

(A) Cumulative budding events occurring prior to and following abrupt glucose withdrawal in a microfluidic chamber, with growth dynamics observed by microscopy.

Prototrophic cells (yLB128) were grown in synthetic media lacking amino acids, with and without glucose. Data are an aggregation of three independent biological replicates.

(B) Distributions of mitochondrial to total cell volume ratio in cells (yLB128) growing in synthetic media without amino acids, before or following the shift to identical media lacking either glucose or nitrogen (ammonium sulfate), performed at time 0 min. N ≥ 360 cells.

(C) Heat map depicting the distribution of mitochondrial sphericity values for cells (yLB126) growing in a microfluidics unit in synthetic media containing high glucose, before and after a nutrient shift to identical media containing high galactose in lieu of glucose, performed at time 0 min. Intensity reflects the number of cells possessing a sphericity index within a given 0.01-sphericity-unit bin. N = 354 cells.

(D) Heat maps of mitochondrial sphericity for yLB126, before and during glucose starvation, with data partitioned by budded or unbudded status at the moment of glucose washout. N ≥ 404 cells.

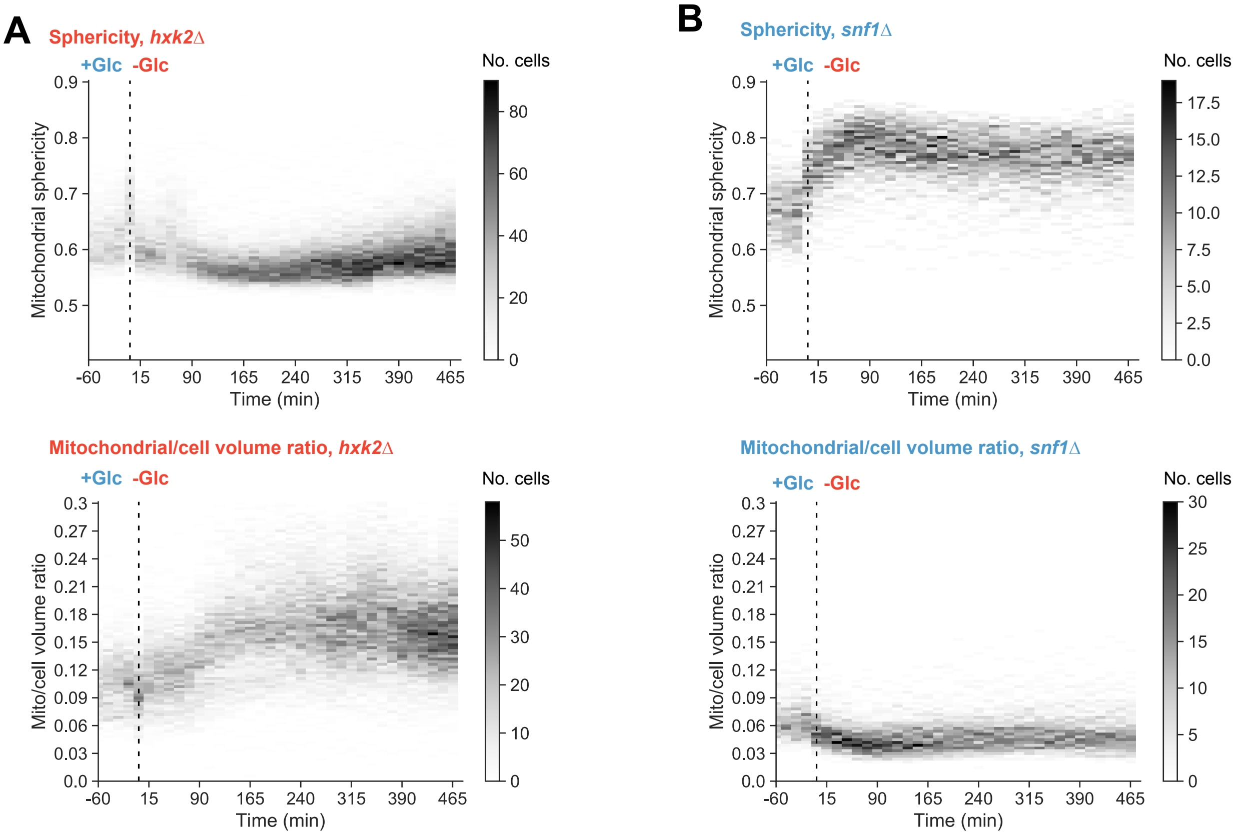


**Figure S4. Disruption of glucose signaling and utilization abrogate bimodal behavior during starvation. Related to Figure 4**

(A) Time-resolved heat maps of mitochondrial-to-total cell volume ratio and sphericity in N = 523 *hxk2Δ* cells (yLB146), before and during acute glucose starvation occurring at 0 min. Compare to Figures 1G and 1H.

(B) Time-resolved heat maps of mitochondrial-to-total cell volume ratio and sphericity in N = 342 *snf1Δ* cells (yLB168), before and during acute glucose starvation.

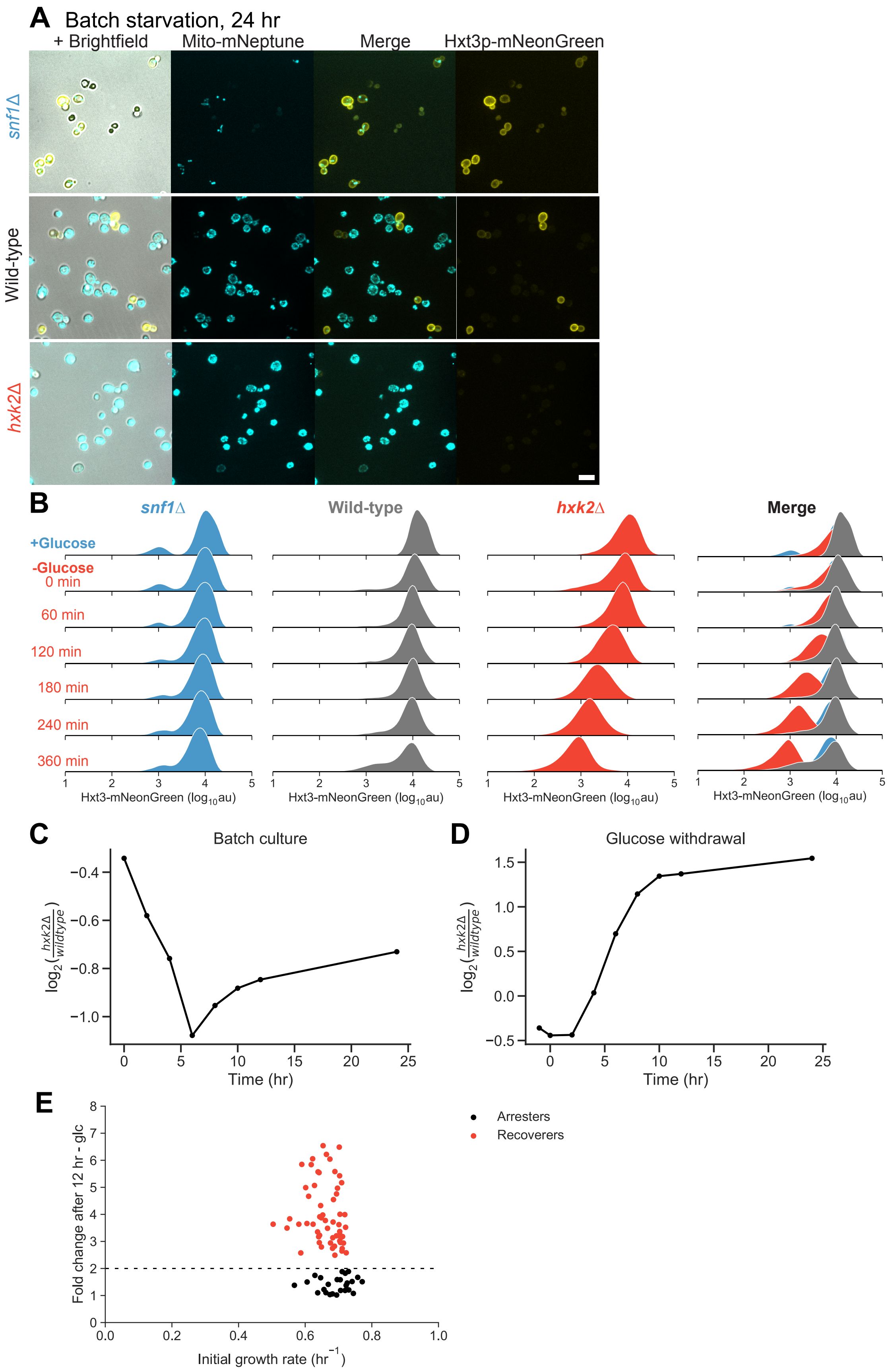
A
Batch starvation, 24 hr
+ Brightfield
Mito-mNeptune
Merge
Hxt3p-mNeonGreen
snf1Δ
Wild-type
hxk2Δ
B
snf1Δ
Wild-type
hxk2Δ
Merge
+Glucose
-Glucose
0 min
60 min
120 min
180 min
240 min
360 min
Hxt3-mNeonGreen (log10au)
C
Batch culture
log2(hxk2Δ/wildtype)
Time (hr)
D
Glucose withdrawal
E
Arresters
Recoverers
Fold change after 12 hr - glc
Initial growth rate (hr-1)

**Figure S5. Fast adaptation to glucose withdrawal confers a reciprocal fitness cost under well-fed condition. Related to Figure 5**

(A) Representative images of cells co-expressing fluorescently labeled Hxt3p-mNeonGreen and mitochondrial matrix-targeted mito-mNeptune in *snf1Δ* (yLB299), *hxk2Δ* (yLB297), or wild-type (yLB256) mutant backgrounds following 24 hr of glucose deprivation. While *hxk2Δ* mutants display uniform mitochondrial network increase and loss of Hxt3p signal and *snf1Δ* uniformly retain Hxt3p-mNeonGreen and have fragmented mitochondria (excepting the shriveled, phase-dark dead cells), the wild-type population is a heterogeneous distribution of Hxt3p-positive cells with mitochondrial fragmentation and Hxt3p-negative cells with large mitochondrial networks. Scale bar, 10 μm.

(B) Hxt3p-mNeonGreen intensity measured by flow cytometry in wild-type, *snf1Δ*, and *hxk2Δ* cells growing in the presence of high glucose and deprived of glucose for the indicated time intervals. Distributions consist of three biological replicates of N = 40,000 cells each.

(C)-(D) Cultures were initiated with equal proportions of wild-type (yLB365) and *hxk2Δ* (yLB373) cells, each expressing distinct fluorescent markers, in synthetic media containing high glucose. The proportions of wild-type and *hxk2Δ* cells were measured during continued cultivation in high glucose *(C)* and following sudden glucose withdrawal *(D)*. N = 40,000 cells measured by flow cytometry at each time point. In (C), the fraction of *hxk2*Δ cells reaches a minimum at the onset of the diauxic shift, which occurs when all glucose has been fermented, and then rises as cells begin respiring ethanol.

(E) Pre-starvation growth rate of microcolonies founded by single cells plotted against the change in the cell lineage’s size following abrupt glucose withdrawal from the media. Lineages are assigned to two states (recoverers (red) and arresters (black) by their success or failure in doubling in size over the first 12 hr of glucose starvation).

### SUPPLEMENTARY MOVIES

**Movie S1. Heterogeneity in mitochondrial morphology during sudden glucose starvation.**

Time lapse imaging of mito-EGFP (yLB113) (right panel), with brightfield overlay (left panel) during growth in synthetic media containing glucose (blue background) and during abrupt glucose washout (red background). Cells display heterogeneity in their ability to retain mitochondrial tubular structure and perform mitochondrial biogenesis. The presence of multiple clonal clusters showing no collapse is atypical, and this field is shown here for the purposes of clearly displaying the differences between divergent phenotypes. Time stamp indicates hours:minutes relative to glucose withdrawal occurring at time 0 hr. Time interval between frames, 45 min. Scale bar, 10 μm.

**Movie S2. Diversity in mitochondrial structural state during acute glucose starvation, then uniform morphology during refeeding.**

Time lapse of cells shown in Figure 1D expressing mito-mNeonGreen (yLB126) (right panel) with brightfield overlay (left panel), imaged during growth in synthetic media containing glucose (blue background), for 8 hr glucose starvation (red background), and for 4 hr during the refeeding on high-glucose media (blue background). Time stamp indicates hours:minutes relative to glucose withdrawal occurring at time 0 hr. Glucose is returned to the media at 8 hr. Time interval between frames, 15 min. Scale bar, 10 μm.

**Movie S3. Mutants in glucose signaling and utilization adopt homogeneous cell fates during sudden nutrient shifts.**

Time lapse imaging of wild-type (yLB126, top row), *hxk2*Δ (yLB146, middle row), and *snf1*Δ (yLB168, bottom row) cells expressing mito-mNeonGreen, with and without brightfield overlay (right and left columns, respectively), during growth in synthetic media containing high glucose and following abrupt washout of glucose from the media (blue and red backgrounds, respectively). Wild-type cells display heterogeneous arrest and recovery, while *hxk2*Δ mutants recover homogeneously and *snf1*Δ mutants arrest homogeneously. Time stamp indicates hours:minutes relative to glucose washout at time 0 hr. Time interval between frames, 15 min. Scale bar, 10 μm.

**Table S1** Transgenic yeast strains used in this study

| Strain name | Genotype description |
|---|---|
| yLB1 | *W303 MATa can1-100 leu2-3,112 his3-11,15 ura3-1 BUD4-S288C RAD5 TRP* |
| yLB41 | *yLB1 his3::pADH1-preSu9-link-mKate2-ADH1term-HIS3 SEC63-EGFP-KAN* |
| yLB45 | *yLB1 his3::pADH1-preSu9-link-mKate2-ADH1term-HIS3 ABP1-EGFP-KAN* |
| yLB69 | *yLB1 his3::pADH1-preSu9-link-mKate2-ADH1term-HIS3 ABP140-mNeonGreen-KAN* |
| yLB73 | *yLB1 $rho^0$* |
| yLB113 | *W303 MATa can1-100 his3::pADH1-preSu9-link-EGFP-ADH1term-HIS3 BUD4-S288C RAD5 TRP* |
| yLB126 | *yLB1 his3::pADH1-preSu9-link-mNeonGreen-ADH1term-HIS3* |
| yLB128 | *can1-100 his3::pADH1-preSu9-link-mNeonGreen-ADH1term-HIS3 BUD4-S288C* |
| yLB134 | *yLB1 his3::pADH1-preSu9-link-mNeonGreen-ADH1term-HIS3 dnm1::KAN* |
| yLB145 | *yLB1 hxk2::NAT* |
| yLB146 | *yLB126 hxk2::NAT* |
| yLB167 | *yLB1 snf1::KAN* |

| | |
|---|---|
| yLB168 | *yLB126 snf1::KAN* |
| yLB180 | *yLB126 mig1::NAT mig2::KAN* |
| yLB181 | *yLB1 mig1::NAT mig2::KAN* |
| yLB194 | *yLB1 reg1::KAN* |
| yLB196 | *yLB126 reg1::KAN* |
| yLB219 | *yLB1 his3::pADH1-preSu9-link-ratiometric pHluorin-HIS3 ura3-1* |
| yLB232 | *yLB1 snf3::KAN rgt2::NAT* |
| yLB233 | *yLB126 snf3::KAN rgt2::NAT* |
| yLB256 | *yLB1 his3::pADH1-preSu9-link-mNeptune-ADH1term-HIS3 HXT3-mNeonGreen-CgLEU2* |
| yLB297 | *yLB256 hxk2::NAT* |
| yLB299 | *yLB256 snf1::KAN* |
| yLB365 | *yLB1 his3::pACT1-mNeptune-ADH1t-HIS3* |
| yLB373 | *yLB1 his3::pACT1-mNeonGreen-ADH1t-HIS3 hxk2::NAT* |
| yLB397 | *yLB1 his3::pACT1-ratiometric pHluorin-ADH1term-HIS3* |
| yLB412 | *yLB397 his3::pACT1-ratiometric pHluorin-ADH1term-HIS3 snf1::KAN* |
| yLB416 | *yLB397 his3::pACT1-ratiometric pHluorin-ADH1term-HIS3 hxk2::NAT* |
| yLB432 | *W303 MATa can1-100 leu2-3,112 his3-11,15 BUD4-S288C RAD5 TRP URA HXT3-mNeonGreen-SpHIS5 HXT7-mNeptune-CgLEU2* |
| yLB453 | *BC187 HIS3::pHXT3-HXT3-link-mNeonGreen-ADH1t-KAN-HIS3* |
| yLB457 | *Y12 HIS3::pHXT3-HXT3-link-mNeonGreen-ADH1t-KAN-HIS3* |
| yLB463 | *YJM978 HIS3::pACT1-mNeptune-ADH1t-KAN-HIS3* |
| yLB467 | *CEN.PK HIS3::pACT1-mNeptune-ADH1t-KAN-HIS3* |
| yLB470 | *DBVPG1373 HIS3::pACT1-mNeptune-ADH1t-KAN-HIS3* |
| yLB474 | *L-1374 HIS3::pACT1-mNeptune-ADH1t-KAN-HIS3* |
| yLB478 | *BC187 HIS3::pACT1-mNeptune-ADH1t-KAN-HIS3* |
| yLB480 | *YS2 HIS3::pACT1-mNeonGreen-ADH1t-KAN-HIS3* |
| yLB486 | *Y12 HIS3::pACT1-mNeptune-ADH1t-KAN-HIS3* |
| yLB492 | *K11 HIS3::pACT1-mNeptune-ADH1t-KAN-HIS3* |
| yLB494 | *YPS606 HIS3::pACT1-mNeptune-ADH1t-KAN-HIS3* |
| yLB496 | *UWOPS83-787.3 HIS3::pACT1-mNeptune-ADH1t-KAN-HIS3* |

**Table S2** Plasmids used in the construction of transgenic yeast strains

| Plasmid | Base vector | Insert | Strains with insert integrated by homologous recombination |
|---|---|---|---|
| pLB36 | pRS403 | pADH1-preSu9-link-mKate2-ADH1t-HIS3 | yLB41, yLB45, yLB69 |
| pLB39 | pRS403 | pADH1-preSu9-link-mNeptune-HIS3 | yLB256, yLB297, yLB299 |
| pLB52 | pUC19 | SEC63-link-yEGFP-ADH1t-KAN-3'SEC63 | yLB41 |
| pLB54 | pUC19 | ABP1-link-yEGFP-ADH1t-KAN-3'ABP1 | yLB45 |

| | | | |
|---|---|---|---|
| pLB57 | pUC19 | ABP140-link-mNeonGreen-ADH1t-KAN-3'ABP140 | yLB69 |
| pLB76 | pRS403 | pADH1-preSu9-link-mNeonGreen-ADH1t-HIS3 | yLB126, yLB146, yLB168, yLB180, yLB196, yLB233 |
| pLB104 | pRS403 | pADH1-preSu9-ratiometric pHluorin-ADH1t-HIS3 | yLB219 |
| pLB114 | pUC19 | HXT3-link-mNeonGreen-ADH1t-CgLEU2-3'HXT3 | yLB256, yLB297, yLB299 |
| pLB142 | pUC19 | HXT7-link-mNeptune-ADH1t-CgLEU2-3'HXT7 | yLB432 |
| pLB144 | pUC19 | HXT3-link-mNeonGreen-ADH1t-SpHIS5-5'HXT3 | yLB432 |
| pLB152 | pRS403 | pACT1-mNeptune-ADH1t-HIS3 | yLB365 |
| pLB153 | pRS403 | pACT1-mNeonGreen-ADH1t-HIS3 | yLB373 |
| pLB161 | pRS403 | pACT1-ratiometric pHluorin-ADH1t-HIS3 | yLB397, yLB412, yLB416 |
| pLB164 | pRS403 | pHXT3-HXT3-link-mNeonGreen-ADH1t-KAN-HIS3 | yLB453, yLB457 |
| pLB166 | pRS403 | pACT1-mNeonGreen-ADH1t-KAN-HIS3 | yLB480 |
| pLB168 | pRS403 | pACT1-mNeptune-ADH1t-KAN-HIS3 | yLB463, yLB467, yLB470, yLB474, yLB478, yLB486, yLB492, yLB494, yLB496 |
